# Targeting Macrophage Connexin 43 Hemichannels Suppresses Synovial Inflammation and Osteoarthritis

**DOI:** 10.64898/2026.09.04.749181

**Authors:** Rui Hua, Yi Tian, Xuewei Wang, Frank A. Buttacavoli, Animesh Agarwal, Liang Ma, Manuel A. Riquelme, Hang Xu, Lidan Zhang, Teja Guda, Jun Zhang, Sumin Gu, Jean X. Jiang

## Abstract

Osteoarthritis (OA) is the most common chronic joint disease, yet no disease-modifying therapies exist. Synovial inflammation is increasingly recognized as a key driver of OA progression, but its therapeutic targets remain poorly defined. Here we show that selective inhibition of connexin 43 (Cx43) hemichannels (HCs) attenuates OA progression in a murine post-traumatic OA model. AAV-mediated Cx43 deletion reduced cartilage degeneration, synovial inflammation, subchondral bone sclerosis, and pain. We developed a monoclonal antibody, Cx43(M1), that selectively blocks Cx43 HCs without disrupting gap junction communication. Cx43(M1) preserved joint integrity and alleviated OA even when treatment began after cartilage damage was established. The antibody preferentially targeted synovial macrophages, suppressing HC activity, ATP and PGE₂ release, and inflammatory gene expression. Macrophage-specific Cx43 deletion reproduced these effects, and Cx43 HC blockade similarly inhibited inflammatory responses in primary human OA synovial macrophages. These findings identify synovial macrophage Cx43 HCs as promising therapeutic targets for OA.

## Introduction

Osteoarthritis (OA) is the most prevalent chronic joint disease and a leading cause of global disability. While historically viewed as a simple “wear-and-tear” degeneration of articular cartilage, accumulating evidence suggests that OA is a complex, whole-joint disease characterized by progressive cartilage erosion, subchondral bone remodeling, synovial inflammation, and persistent pain [1]. Despite its massive clinical burden, there are currently no approved disease-modifying therapies for OA. Recent studies have increasingly recognized synovium as a critical contributor to OA pathogenesis [2–4]. During disease progression, synovial macrophages and fibroblasts transition into a pro-inflammatory state, secreting potent cytokines such as IL-1β, TNF-α, and IL-6, as well as catabolic enzymes like matrix metalloproteinases (MMPs) and aggrecanases (ADAMTS) [5, 6]. These mediators amplify local inflammation and stimulate chondrocytes to produce matrix-degrading enzymes, thereby accelerating cartilage destruction. Importantly, chronic synovitis is now recognized not merely as a secondary consequence of cartilage injury but as an early and persistent driver of OA progression, strongly associated with structural deterioration and pain severity. However, the upstream molecular mechanisms governing inflammatory activation within the synovium remain poorly defined, limiting the development of effective disease-modifying strategies.

Connexin 43 (Cx43) is a membrane protein with four transmembrane domains critical for maintaining musculoskeletal homeostasis. Cx43 forms both gap junctions and hemichannels (HCs), which permit the passage of small molecules (< 1.2 kDa). Cx43 gap junctions enable intercellular communication between adjacent cells. In contrast, Cx43 HCs, which are unpaired gap junction channels, mediate communication between the intracellular and extracellular environments [7]. Under physiological conditions, Cx43 HCs typically remain closed. However, they can be activated by mechanical stress or pro-inflammatory cytokines to release potent signaling molecules, most notably adenosine triphosphate (ATP) and prostaglandin E_2_ (PGE_2_) [8–10]. Emerging evidence suggests that Cx43 HCs are highly active in inflammatory settings, where they regulate ATP-dependent purinergic signaling and inflammasome activation [11]. Furthermore, PGE₂ contributes to pain sensitization by acting on nociceptive pathways and enhances the production of pro-inflammatory mediators, thereby creating a positive feedback loop that exacerbates joint pathology [12, 13].

Recent studies have implicated Cx43 in OA pathogenesis [14]. Increased Cx43 expression has been observed in joint tissues of OA patients, including cartilage and synovium [15, 16], where it promotes the expression of catabolic and pro-inflammatory mediators such as MMPs, ADAMTS enzymes, and cytokines [17]. In addition, Cx43-mediated ATP release has been associated with chondrocyte dysfunction and cartilage degeneration in arthritic joints [18]. However, most prior studies have focused on total Cx43 expression or gap junction-mediated signaling, leaving the specific contribution of Cx43 HCs unresolved. Moreover, whether Cx43 HCs within synovial macrophages function as upstream regulators of inflammatory signaling and OA progression remains unknown. Addressing this gap is particularly important because macrophages are increasingly recognized as central orchestrators of synovial inflammation and joint degeneration.

In this study, we investigated the therapeutic potential of targeting Cx43 HCs to attenuate OA progression. Using AAV-mediated genetic knockdown and a novel monoclonal antibody, Cx43(M1), we demonstrate that inhibition of Cx43 HC activity attenuates synovial inflammation, pain-related behaviors, and joint structural deterioration in a murine OA model. We further identify synovial macrophages as a major cellular target of Cx43-mediated signaling and show that macrophage-specific deletion of Cx43 recapitulates the protective effects of HC blockade. Mechanistically, we demonstrate that Cx43 regulates ATP and PGE_2_ release and downstream inflammatory gene expression across multiple macrophage populations, including primary human cells. Collectively, these findings identify synovial macrophage Cx43 HCs as a key upstream driver of OA and shift the field beyond a predominantly chondrocyte-centered model toward inflammation-directed therapeutic intervention.

## Results

### Knocking down Cx43 expression in joint tissues alleviates surgery-induced OA

To determine whether Cx43 deletion in joint tissue could ameliorate OA symptoms, we used an adeno-associated virus (AAV) mediated Cre recombinase approach in Cx43 flox/flox mice using post-traumatic OA model induced by surgical destabilization of the medial meniscus (DMM). AAV5 expressing GFP or Cre-GFP was delivered by intra-articular injection into OA knee joints. Two months post-injection, AAV expression remained detectable in multiple joint tissues, including articular cartilage and synovium (Fig. 1A and Fig. S1), demonstrating sustained gene expression in the knee joint. Intra-articular injection of AAV-Cre effectively reduced Cx43 level in both synovium and cartilage, with the greatest reduction observed in synovial tissue (Fig. 1B, C). As illustrated in Fig. 1D, AAV was administered intra-articularly 7 days before sham or DMM surgery, and joint tissues were collected 8 weeks after surgery. Cx43 deletion attenuated proteoglycan loss and cartilage erosion, as demonstrated by Safranin O/Fast Green staining (Fig. 1E). Osteoarthritis Research Society International (OARSI) scoring confirmed preservation of articular cartilage structure, with no significant differences between AAV-control and Cre-injected sham groups (Fig. 1F). H&E staining further showed reduced synovial hyperplasia following Cx43 deletion (Fig. 1G, H).

**Figure 1.**
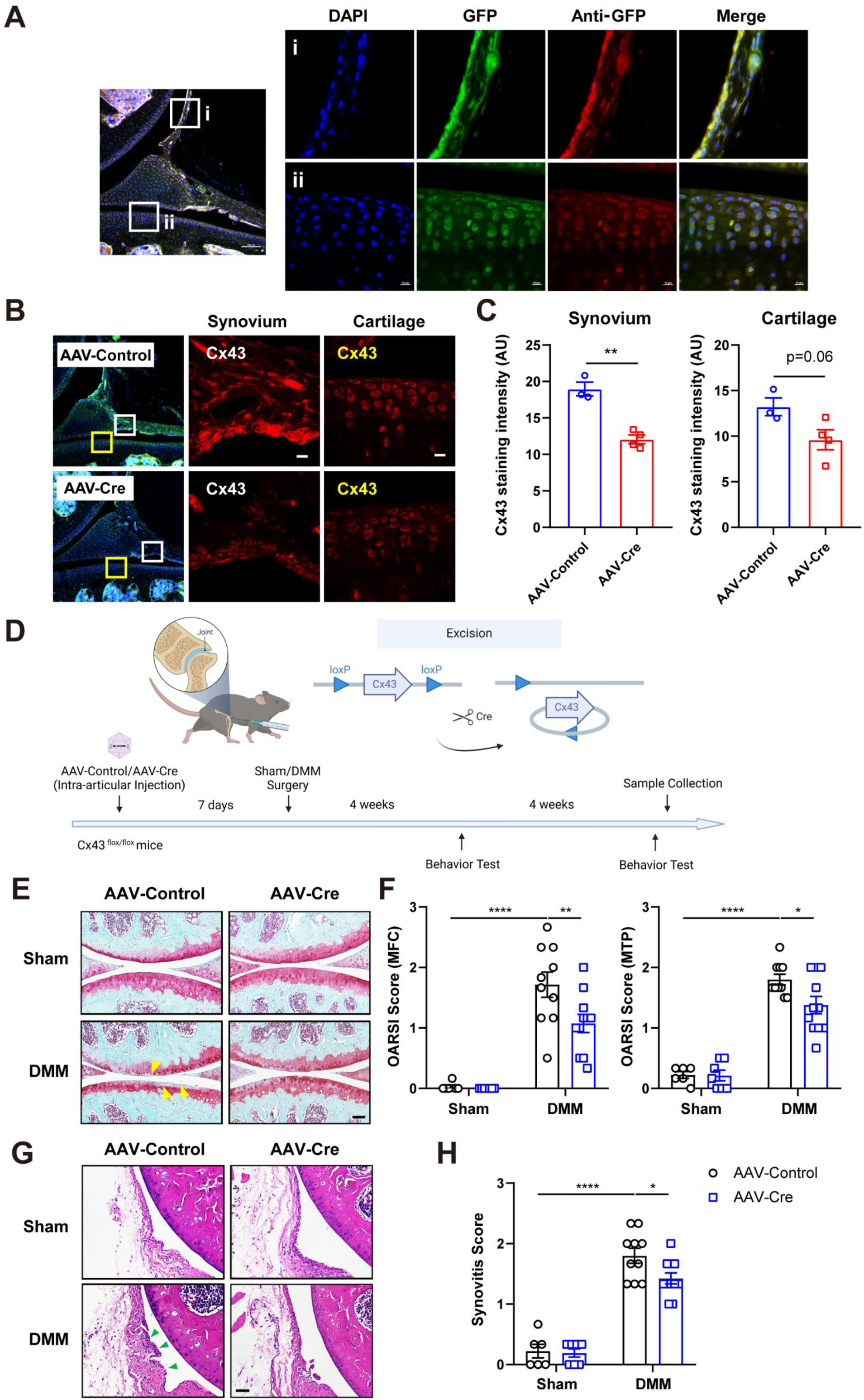
Knocking down Cx43 expression attenuates articular cartilage breakdown and reduced synovium inflammation in the DMM model. **(A)** Representative images of AAV-GFP expression in the synovium (i, upper panels) and articular cartilage (ii, lower panels) regions 2 months after intra-articular injection. Expression was visualized by direct GFP fluorescence (green) or by anti-GFP immunostaining (red) in cryosections collected. Scale bar, 10 μm. **(B)** Representative images of Cx43 immunofluorescence staining in Cx43 flox/flox mice injected with AAV-Control (upper panels) or AAV-Cre (lower panels). Synovium (white box) and articular cartilage (yellow box) regions are shown in higher magnification. Scale bar, 10 μm. **(C)** Quantification of Cx43 expression levels in the synovium (left panel) and cartilage (right panel), respectively. Staining intensity was quantified using NIH ImageJ software. n = 3-4 mice/group. **(D)** Schematic diagram of the timeline for surgical procedures, virus injection, and outcome assessments. AAV was injected 7 days before sham or DMM surgery. Behavioral tests were conducted every 4 weeks, and tissue samples were collected 8 weeks after surgery. **(E)** Representative images of Safranin O/Fast Green staining in AAV-Control or AAV-Cre-injected Cx43 flox/flox mice after sham or DMM surgery. Areas of cartilage destruction are indicated by yellow arrowheads. Scale bar, 200 μm. **(F)** Averaged OARSI scores for the medial femoral condyle (MFC, left panel) and medial tibial plateau (MTP, right panel). **(G,H)** Representative images of H&E staining **(G)** and quantification of synovitis score **(H)** in AAV-Control or AAV-Cre-injected Cx43 flox/flox mice after sham or DMM surgery. Areas of synovium inflammation are indicated by green arrowheads. Scale bar, 200 μm. n = 6-11 mice/group. Data are presented as mean ± SEM. *, p < 0.05; **, p < 0.01; ****, p < 0.0001.

Since OA is characterized by abnormal subchondral bone remodeling [19], we analyzed subchondral bone microarchitecture using micro-computed tomography (micro-CT). Eight weeks after DMM surgery, bone volume/total volume (BV/TV), trabecular thickness (Tb.Th), and bone mineral density (BMD) were significantly increased in the medial tibial subchondral bone compared with sham controls (Fig. 2A-D). Deletion of Cx43 significantly reduced BV/TV, Tb.Th, and BMD (Fig. 2B-D), indicating attenuation of OA-associated subchondral bone sclerosis. OA progression is also accompanied by prominent pain-related behaviors. To evaluate mechanical hypersensitivity, paw withdrawal thresholds were measured using von Frey filaments, and spontaneous locomotor activity was assessed by the open field test. At both 4 and 8 weeks following DMM surgery, AAV-Cre treated Cx43 flox/flox mice exhibited significantly higher paw withdrawal thresholds than AAV-Control mice (Fig. 2E, H), indicating reduced mechanical allodynia. Furthermore, Cx43 deletion in joint tissues increased total distance traveled and decreased immobility time in the open field assay (Fig. 2F, G, I, J), reflecting improved spontaneous activity. Together, these findings suggest that Cx43 deletion in joint tissues alleviates OA-associated structural degeneration, reduces synovial hyperplasia, and improves pain-related behaviors and functional impairment.

**Figure 2.**
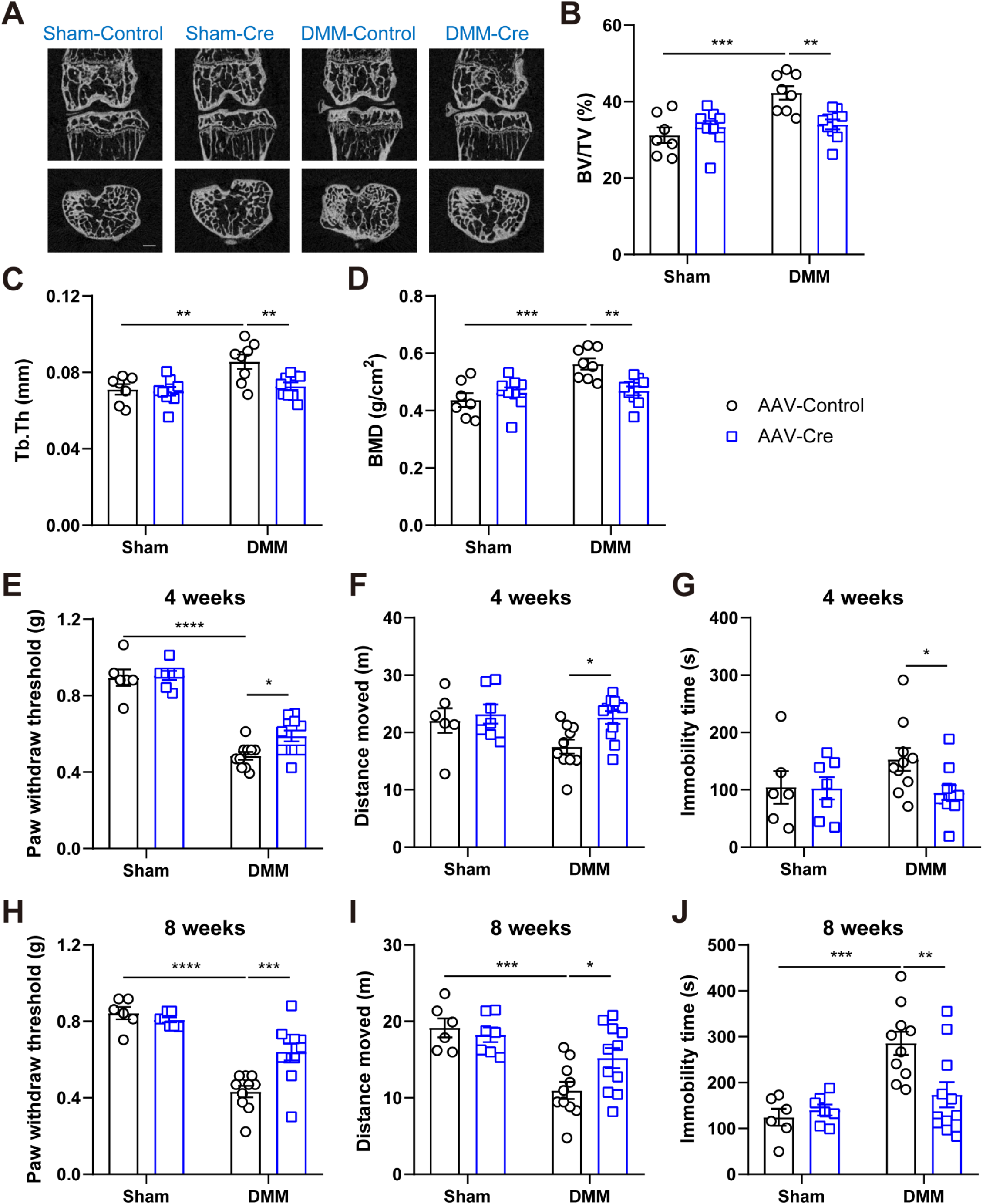
AAV-mediated Cx43 knockdown protects against subchondral bone sclerosis and OA-related pain behaviors in the DMM model. **(A)** Representative two-dimensional micro-CT images of the medial and lateral subchondral bone compartments of the tibial plateau 8 weeks after surgery. **(B-D)** Quantification of subchondral trabecular bone parameters, including bone volume fraction (BV/TV), trabecular thickness (Tb.Th), and bone mineral density (BMD). n = 7-9 mice/group. **(E, H)** Hind paw withdrawal thresholds were measured using the von Frey filament test 4 weeks **(E)** and 8 weeks **(H)** post-surgery. **(F,G, I,J)** Open-field behavioral assessments were conducted 4 weeks **(F,G)** and 8 weeks **(I,J)** post-surgery. n = 6-11 mice/group. Data are presented as mean ± SEM. *, p < 0.05; **, p < 0.01; ***, p < 0.001; ****, p < 0.0001. Abbreviations: TV, total volume; BV, bone volume; Tb.Th, trabecular thickness.

### Inhibition of Cx43 hemichannel ameliorates OA progression

To specifically target Cx43 HCs, we used our previously developed monoclonal antibody, Cx43(M1), which selectively blocks HC activity without disrupting gap junction communication [20, 21]. Cx43(M1) antibody (25 mg/kg) was administered intraperitoneally 30 minutes and 4 weeks after sham or DMM surgery, followed by tissue collection at 8 weeks as illustrated in Fig. 3A. Safranin O/Fast Green staining showed that Cx43(M1) treatment preserved articular cartilage integrity and proteoglycan content compared with the vehicle group, resulting in significantly lower OARSI scores (Fig. 3B, C). In addition, Cx43(M1) substantially reduced synovitis severity in DMM-operated mice, as reflected by decreased synovitis scores (Fig. 3D, E). Collagen X (Col X) is a marker of hypertrophic chondrocytes and matrix metalloproteinase 13 (MMP13), a major enzyme responsible for degradation of type II collagen in articular cartilage [22, 23], were both elevated following DMM surgery. Immunohistochemical staining of articular cartilage revealed that Cx43(M1) significantly suppressed OA-induced increase of Col X and MMP13 expression within articular cartilage (Fig. 3F, G). Moreover, MMP13 expression in synovial cells was reduced in the Cx43(M1)-treated joint compared to the vehicle group (Fig. 3H, I), suggesting that broader suppression of catabolic activity within the joint microenvironment. These findings demonstrate that selective inhibition of Cx43 HCs attenuates OA-associated cartilage degeneration and synovial pathology while reducing markers of chondrocyte hypertrophy and matrix catabolism.

**Figure 3.**
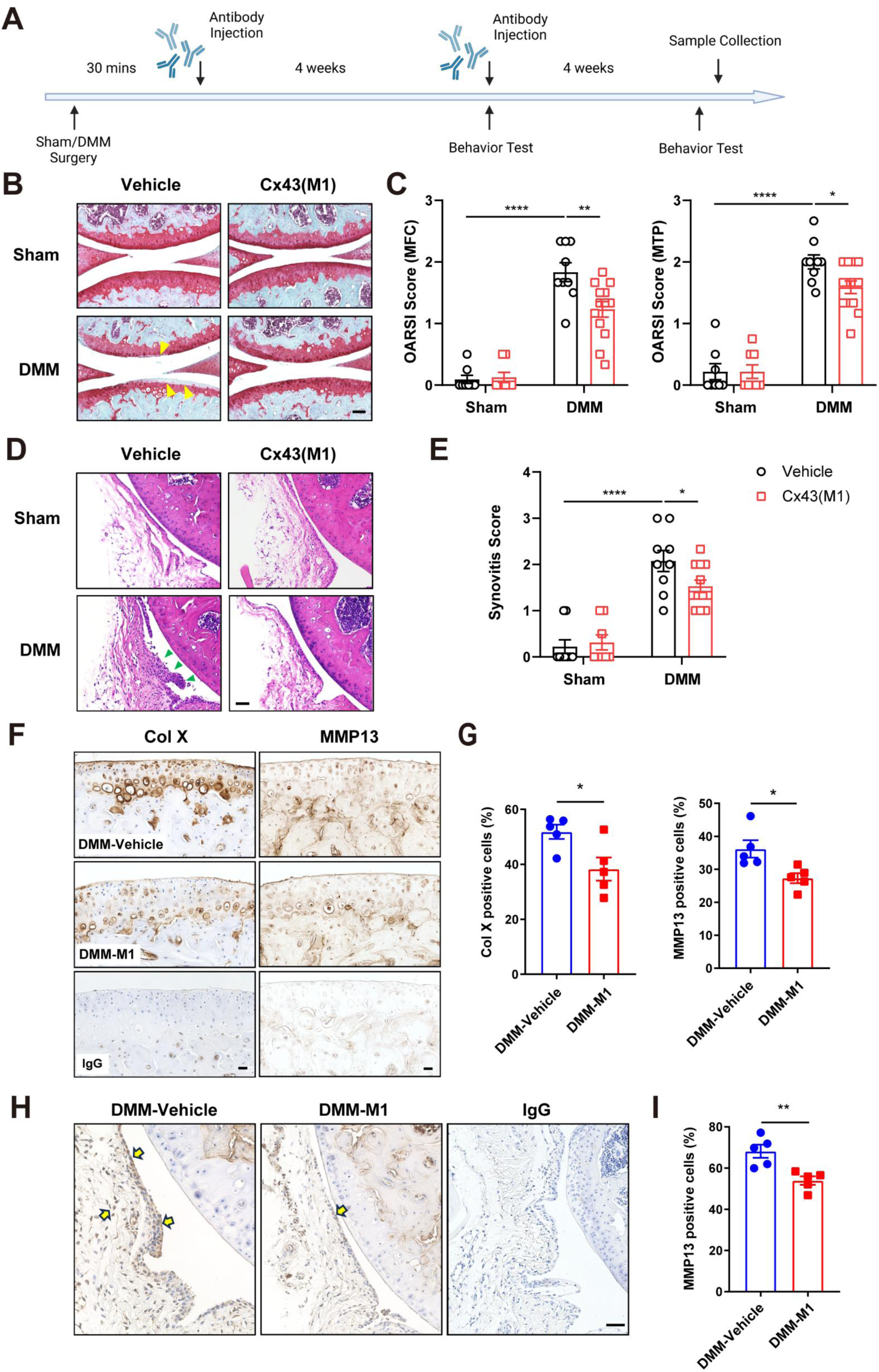
Cx43 antibody treatment preserves cartilage integrity and reduces synovial inflammation, accompanied by decreased Col X and MMP13 expression in the DMM model. **(A)** Schematic diagram of the timeline for surgical procedures, antibody injections, and outcome assessments. Cx43 hemichannel-blocking antibody was administered 30 minutes and 4 weeks after surgery. Behavioral tests were conducted every 4 weeks, and tissue samples were collected 8 weeks after surgery. **(B)** Representative images of Safranin O/Fast Green staining in vehicle or Cx43(M1) antibody-treated mice after sham or DMM surgery. Areas of cartilage destruction are indicated by yellow arrowheads. Scale bar, 200 μm. **(C)** Averaged OARSI scores for medial femoral condyle (MFC, left panel) and medial tibial plateau (MTP, right panel). **(D,E)** Representative images of H&E staining **(D)** and quantification of synovitis scores **(E)** in vehicle or Cx43(M1) antibody-treated mice after sham or DMM surgery. Areas of synovial inflammation are indicated by green arrowheads. Scale bar, 200 μm. n = 8-13 mice/group. **(F)** Representative images of immunohistochemical staining for Col X (left panels) and MMP13 (right panels) in articular cartilage of mice subjected to DMM surgery and treated with vehicle or Cx43(M1) antibody. Mouse or rabbit IgG was used as an isotype control (lower panels). Scale bar, 50 μm. **(G)** Quantification of the percentage of Col X-positive (left panel) and MMP13-positive (right panel) chondrocytes. **(H)** Representative images of immunohistochemical staining for MMP13 in synovium. Scale bar, 100 μm. **(I)** Quantification of the percentage of MMP13-positive synoviocytes. n = 5 mice/group. Data are presented as mean ± SEM. *, p < 0.05; **, p < 0.01; ****, p < 0.0001.

Micro-CT analysis of subchondral bone revealed that blocking Cx43 HCs decreased BV/TV and Tb.Th in DMM-induced OA mice, with a trend toward reduced BMD (Fig. 4A-D). In contrast, Cx43(M1) treatment did not alter subchondral bone parameters in sham-operated mice (Fig. 4A-D), suggesting that HC inhibition selectively mitigates pathological bone remodeling associated with OA rather than affecting normal bone homeostasis. OA-associated pain behaviors were evaluated 4 and 8 weeks after surgery. At 4 weeks, Cx43(M1)-treated DMM mice showed trends toward increased von Frey hind paw withdrawal thresholds and greater total distance traveled in the open field test, along with reduced immobility time compared with vehicle-treated mice (Fig. 4E-G). By 8 weeks post-surgery, Cx43(M1) treatment significantly increased paw withdrawal thresholds and improved spontaneous locomotor activity (Fig. 4H-J), indicating progressive improvement in OA-associated pain and functional impairment.

**Figure 4.**
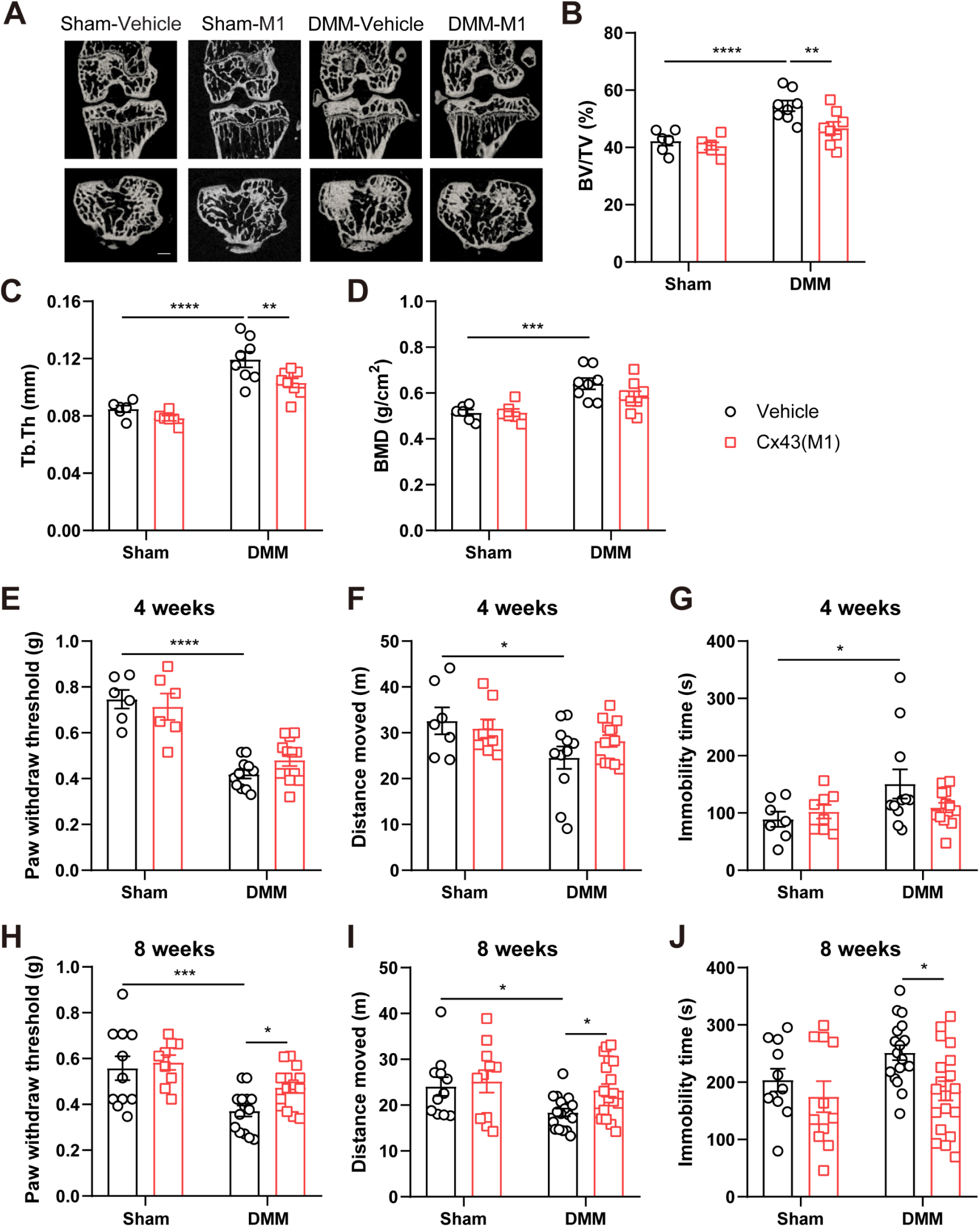
Inhibition of Cx43 hemichannels mitigates subchondral bone sclerosis and OA-associated pain symptoms in the DMM model. **(A)** Representative two-dimensional micro-CT images of the medial and lateral subchondral bone compartments of the tibial plateau 8 weeks after surgery. **(B-D)** Quantification of subchondral trabecular bone parameters, including bone volume fraction (BV/TV), trabecular thickness (Tb.Th), and bone mineral density (BMD). n = 6-8 mice/group. **(E, H)** Hind paw withdrawal thresholds were measured using the von Frey filament test 4 weeks **(E)** and 8 weeks **(H)** post-surgery. **(F,G, I,J)** Open field behavioral assessments were conducted 4 weeks **(F,G)** and 8 weeks **(I,J)** post-surgery. n = 6-13 mice/group. Data are presented as mean ± SEM. *, p < 0.05; **, p < 0.01; ***, p < 0.001; ****, p < 0.0001.

We next investigated whether a single administration of Cx43(M1) could provide sustained protection against OA progression. To test this, mice received a single intraperitoneal injection of Cx43(M1) or saline 30 minutes post-surgery, followed by behavior tests and histological analysis 8 weeks after DMM surgery (Fig. S2A). A single Cx43(M1) significantly reduced the OARSI score in the medial femoral condyle (MFC), while a trend toward reduced cartilage damage was observed in the medial tibial plateau (Fig. S2B, C). Compared to saline-treated controls, mice receiving Cx43(M1) showed a modest increase in paw withdrawal thresholds in response to mechanical stimuli (Fig. S2D). In addition, Cx43(M1)-treated mice exhibited a trend toward increased total distance traveled and reduced immobility time in the open field assay (p = 0.09) (Fig. S2E, F). These findings suggest that inhibition of Cx43 HCs not only attenuates OA-associated structural degeneration and pain-related behaviors but also confers durable therapeutic benefit following a single administration, supporting the potential for sustained therapeutic benefit.

To further evaluate the therapeutic potential of Cx43 HC blockade under conditions more closely resembling clinical intervention, the initial administration of Cx43(M1) was delayed until 10 days after DMM surgery, when cartilage degeneration was already underway [24, 25]. Additional antibody injections were administered on days 25 and 40, and joint tissues were collected 8 weeks after DMM surgery (Fig. S3A). Delayed Cx43(M1) treatment preserved cartilage integrity, as shown by significantly lower OARSI scores compared to vehicle-treated controls (Fig. S3B, C). In addition, delayed antibody administration reduced synovial inflammation, resulting in significantly lower synovitis scores (Fig. S3D, E). These findings demonstrate that Cx43 HC inhibition remains effective even after OA pathology has been established, supporting a therapeutically relevant intervention window for disease attenuation.

### Cx43 hemichannel inhibition suppresses synovial macrophage activation and inflammatory gene expression

Because of the dense extracellular matrix of cartilage and the large molecular size of antibodies, penetration of systemically administered antibody into cartilage is generally limited [26]. To determine the distribution pattern of Cx43(M1) in the knee joint, Cx43 antibody (25 mg/kg) containing a human IgG epitope or saline was administered after DMM surgery. The mouse hindlimbs were collected 1 or 2 weeks after injection and sections were immunolabeled with Alexa Fluor 594-conjugated anti-human IgG secondary antibody. The antibody signal was detectable at both times and was primarily localized to the synovial region (Fig. 5A). To assess Cx43 HC activity *in vivo*, an Evans Blue dye uptake assay was performed. Mice were treated with saline or Cx43(M1) after DMM surgery, and Evans Blue dye was administered two weeks later. DMM increased HC opening by approximately two-fold, which was inhibited by Cx43(M1) treatment (Fig. 5B, C). Immunolabeling further confirmed that the antibody blocked OA-induced HC opening in Cx43-expressing synoviocytes (Fig. S4A). We then co-immunolabeled sections with the macrophage marker CD68 and found co-localization of Cx43(M1) with CD68 in the synovium (Fig. 5D, E). Given the marked synovial hyperplasia and macrophage infiltration observed under OA condition, we further characterized macrophage polarization in the DMM model. M1-like macrophages (pro-inflammatory) were identified by iNOS expression. CD68^+^iNOS^+^ double-positive cells were identified predominantly localized in the synovial lining layer (Fig. S4B). In contrast, the M2-like macrophage (anti-inflammatory) marker showed minimal co-localization with CD68 (Fig. S4C), indicating preferential accumulation of pro-inflammatory M1-like macrophages during OA progression. To determine whether Cx43 HC inhibition modulates macrophage activation, we quantified the percentage of CD68⁺iNOS⁺ double-positive cells in the synovium of the Cx43(M1) treatment. Compared with sham-operated controls, DMM induced an approximately two-fold increase in CD68⁺iNOS⁺ macrophages, which was significantly attenuated by Cx43(M1) treatment (Fig. 5F, G). These findings demonstrate that Cx43 HC blockade preferentially targets synovial macrophages, suppresses OA-associated HC activation, and limits the accumulation of pro-inflammatory macrophages within the synovium.

**Figure 5.**
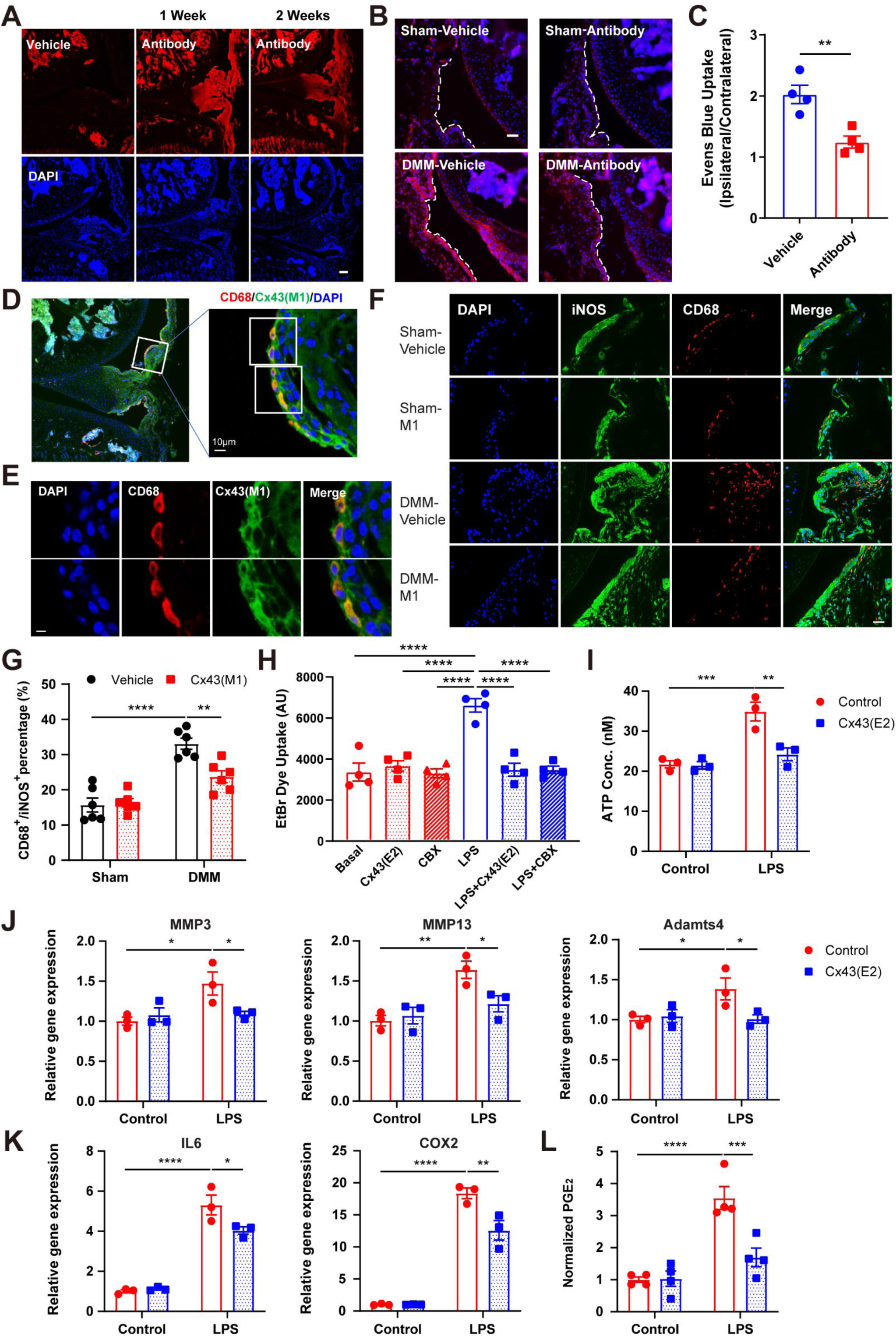
Cx43 hemichannel-blocking antibody targets synovial macrophages to inhibit hemichannel activity and suppress inflammatory responses. **(A)** Cx43 antibody (25 mg/kg) or vehicle was administered intraperitoneally (i.p.) 30 mins after DMM surgery. At 1 or 2 weeks after injection, mice were euthanized and perfused before hindlimb collection. Cryosections were immunolabeled with Alexa Fluor 594-conjugated anti-human IgG secondary antibody. Scale Bar, 100 µm. **(B)** Evans Blue dye (200 mg/kg) was administered by tail vein injection, and hemichannel dye uptake was evaluated by fluorescence microscopy. **(C)** Quantification of Evans Blue fluorescence intensity in the synovium, expressed as a ratio relative to the indicated reference/region, using NIH ImageJ. n = 4 mice/group. **(D,E)** Tissue sections were immunolabeled for the macrophage marker CD68, with two synovial regions shown at higher magnification in **(E)**. The Cx43 antibody co-localized with CD68⁺ macrophages in the synovium. Scale bar, 5 µm. **(F)** Pro-inflammatory M1-like macrophages were identified by co-immunofluorescence staining for CD68 (red) and iNOS (green). **(G)** Quantification of the percentage of CD68^+^iNOS^+^ double-positive cells in vehicle or Cx43(M1) antibody-treated mice after sham or DMM surgery. n = 6 mice/group. **(H)** Synovial macrophages isolated from human OA patients were treated with LPS and either Cx43(E2) antibody or CBX. EtBr dye uptake was quantified by fluorescence microscopy and NIH ImageJ. **(I)** Extracellular ATP release was measured in human OA synovial macrophages treated with LPS and Cx43(E2) antibody. **(J-K)** RNA was extracted and subjected to qRT-PCR analysis for MMP3, MMP13, ADAMTS4, IL6, and COX2 mRNA expression. **(L)** Extracellular PGE_2_ release was measured in human OA synovial macrophages treated with LPS and Cx43(E2) antibody. n = 3-4/group. Data are presented as mean ± SEM. *, p < 0.05; **, p < 0.01; ***, p < 0.001; ****, p < 0.0001.

To establish the role of Cx43 HCs in macrophage inflammatory responses, we utilized the murine macrophage cell line RAW264.7. To investigate the role of Cx43 HCs *in vitro*, we utilized Cx43(E2) antibody, a polyclonal version of Cx43(M1) [27]. EtBr dye uptake assays demonstrated that lipopolysaccharide (LPS) stimulation induced HC opening, which was markedly attenuated by treatment with Cx43(E2) or carbenoxolone (CBX), a widely used gap junction/HC inhibitor [28, 29] (Fig. S5B). Inhibition of Cx43 HCs by Cx43(E2) also suppressed LPS-induced expression of catabolic and inflammatory genes, including MMP3, MMP13, ADAMTS4, IL6, and COX2 (Fig. S5C, D), accompanied by reduced production of the inflammatory mediator PGE_2_ (Fig. S5E). To establish the translational relevance of these findings, we next isolated primary human synovial macrophages from Kellgren-Lawrence (KL) grade 4 knee OA patients. The purity of the isolated macrophages was confirmed by CD68 immunostaining (Fig. S5A). Consistent with the findings in RAW264.7 cells, LPS stimulation induced HC opening in human synovial macrophages, as determined by EtBr dye uptake assays, and this effect was significantly attenuated by Cx43(E2) or CBX treatment (Fig. 5H). Inhibition of Cx43 HCs also reduced LPS-induced extracellular ATP release (Fig. 5I), indicating that Cx43 HCs mediate ATP efflux in synovial macrophages. Furthermore, qRT-PCR analysis demonstrated that Cx43(E2) treatment suppressed LPS-induced upregulation of catabolic and inflammatory genes, including MMP3, MMP13, ADAMTS4, IL6, and COX2 (Fig. 5J, K). Consistent with reduced COX2 expression, inhibition of Cx43 HCs also decreased PGE_2_ production (Fig. 5L). Collectively, these findings demonstrate that Cx43 HCs contribute to inflammatory activation and catabolic responses in both murine and human macrophages.

To determine whether Cx43 HC inhibition also affects synovial fibroblasts, synovial tissue sections were immunolabeled for the fibroblast marker periostin, confirming the presence and fibroblast localization within the synovium (Fig. S6A). In the human synovial fibroblast cell line SW982, stimulation with interleukin-1 beta (IL-1β) significantly increased HC activity, which was attenuated by treatment with either the Cx43(E2) antibody or CBX (Fig. S6B). Furthermore, inhibition of Cx43 HCs suppressed IL-1β-induced expression of MMP3, MMP13, ADAMTS4, ADAMTS5, NOS2, and COX2 (Fig. S6C). These findings demonstrate that Cx43 HCs regulate inflammatory and catabolic responses in synovial macrophages and fibroblasts across both human and murine systems, supporting a conserved role for HC signaling in OA-associated synovial activation.

### Macrophage-specific Cx43 deletion mitigates OA progression

Our findings identified macrophages as a major cellular target of Cx43-mediated signaling within the synovium. To determine the contribution of macrophage-intrinsic Cx43 to OA pathogenesis, we generated a macrophage-specific Cx43 conditional knockout (cKO) mouse model using Lysm-cre and evaluated in the DMM model. Macrophage-specific deletion of Cx43 markedly attenuated OA progression in DMM-operated mice. Safranin O/Fast Green staining revealed reduced cartilage destruction in Cx43 cKO mice compared with Cx43 flox/flox controls, reflected by significantly lower OARSI scores at both the MFC and MTP (Fig. 6A, B). Synovial inflammation was also reduced, as shown by H&E staining and synovitis scoring (Fig. 6C). Micro-CT imaging further showed improved subchondral bone structure in Cx43 cKO mice, with decreased BV/TV, Tb.Th, and BMD in the medial tibial compartments compared with control mice (Fig. 6D-G). Functional assessment of pain-related behaviors indicated no baseline differences between Cx43 flox/flox and cKO mice (Fig. 6H-J). However, following DMM surgery, cKO mice exhibited significantly higher hind paw withdrawal thresholds in the von Frey test at both 4 and 8 weeks (Fig. 6K, N). In the open-field assay, cKO mice showed a trend toward increased distance traveled and reduced immobility 4 weeks post-surgery (Fig. 6L, M), with both measures reaching significance by 8 weeks (Fig. 6O, P). Collectively, these findings demonstrate that macrophage-specific deletion of Cx43 attenuates OA-associated cartilage degeneration, synovial inflammation, subchondral bone sclerosis, and pain-related behavior impairment in the DMM model.

**Figure 6.**
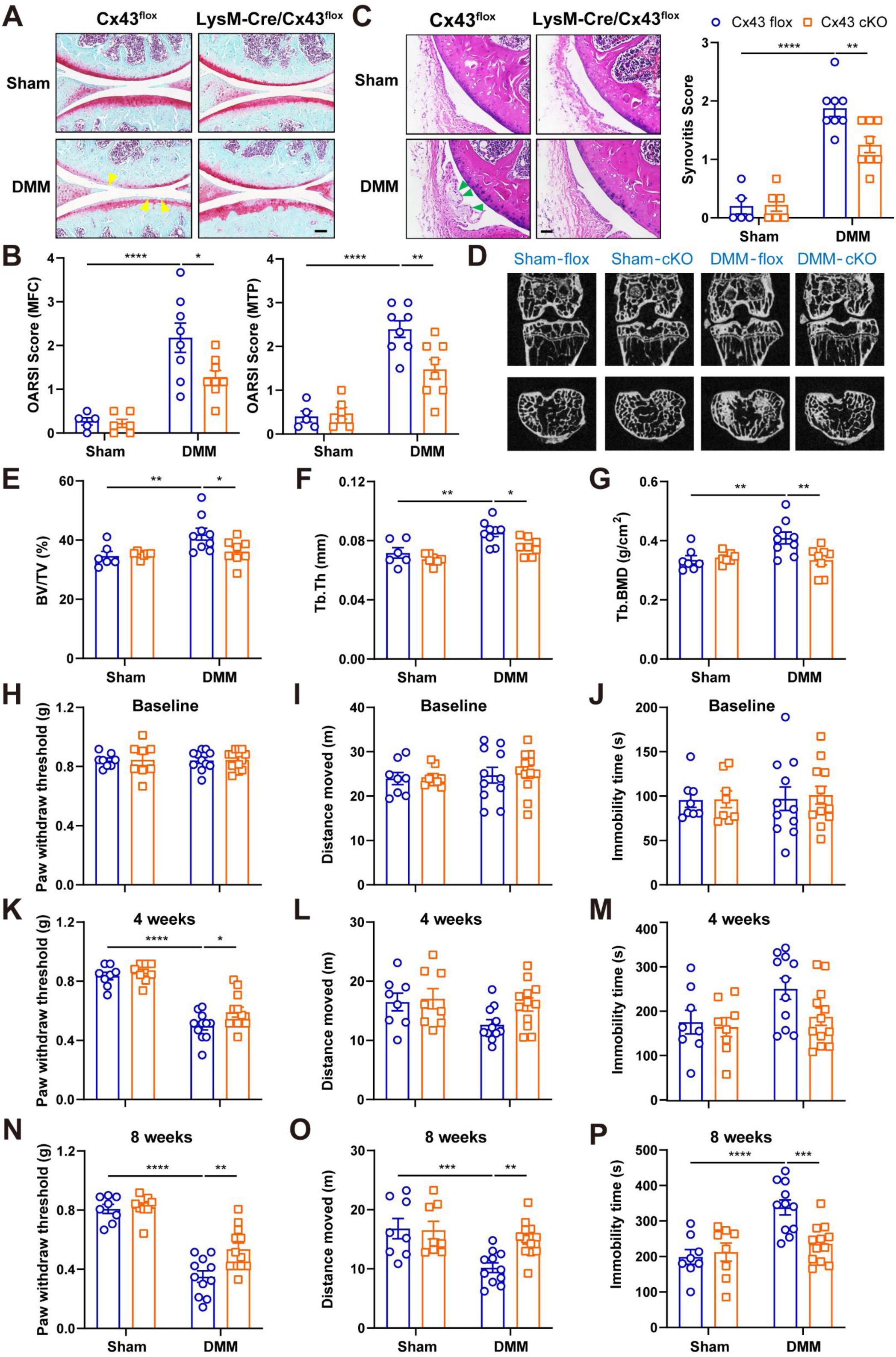
Macrophage-specific Cx43 conditional knockout ameliorates OA progression. **(A)** Representative images of Safranin O/Fast Green histological staining in Cx43 flox/flox or macrophage-specific Cx43 conditional knockout (cKO) mice after sham or DMM surgery. Areas of cartilage destruction are indicated by yellow arrowheads. Scale bar, 200 μm. **(B)** Averaged OARSI scores of medial femoral condyle (MFC, left panel) and medial tibial plateau (MTP, right panel). **(C)** Representative images (left panels) of H&E staining and quantification of synovitis scores (right panel) in Cx43 flox/flox or Cx43 cKO mice after sham or DMM surgery. Areas of synovium inflammation are indicated by green arrowheads. Scale bar, 200 μm. n = 5-8 mice/group. **(D)** Representative two-dimensional micro-CT images of the medial and lateral subchondral bone compartments of the tibial plateau 8 weeks after surgery. **(E-G)** Quantification of subchondral trabecular bone parameters, including bone volume fraction (BV/TV), trabecular thickness (Tb.Th), and bone mineral density (BMD) of subchondral trabecular bone. n = 6-9 mice/group. **(H, K, N)** Hind paw withdrawal thresholds were measured using the von Frey filament test before surgery **(H)** and 4 weeks **(K)** and 8 weeks **(N)** post-surgery. **(I-P)** Open-field behavioral assessments were conducted before surgery **(I,J)** and 4 weeks **(L,M)** and 8 weeks **(O,P)** post-surgery. n = 8-12 mice/group. Data are presented as mean ± SEM. *, p < 0.05; **, p < 0.01; ***, p < 0.001; ****, p < 0.0001.

Having established that macrophage-specific deletion of Cx43 attenuates OA progression *in vivo*, we next investigated the underlying cellular mechanisms in macrophages. Bone marrow-derived macrophages (BMDMs) isolated from Cx43 cKO mice exhibited markedly reduced Cx43 expression compared with Cx43 flox/flox controls (Fig. 7A). Stimulation with LPS induced HC opening, as measured by EtBr dye uptake, which was attenuated in BMDMs isolated from Cx43 cKO mice (Fig. 7B). Consistently, LPS-induced extracellular ATP release was significantly reduced in Cx43-deficient BMDMs (Fig. 7C). At the transcriptional level, genetic deletion of Cx43 suppressed LPS-induced upregulation of ADAMTS4, ADAMTS5, MMP13, TNFα, IL6, and COX2 (Fig. 7D, E). This was accompanied by a corresponding reduction in PGE₂ production (Fig. 7F).

**Figure 7.**
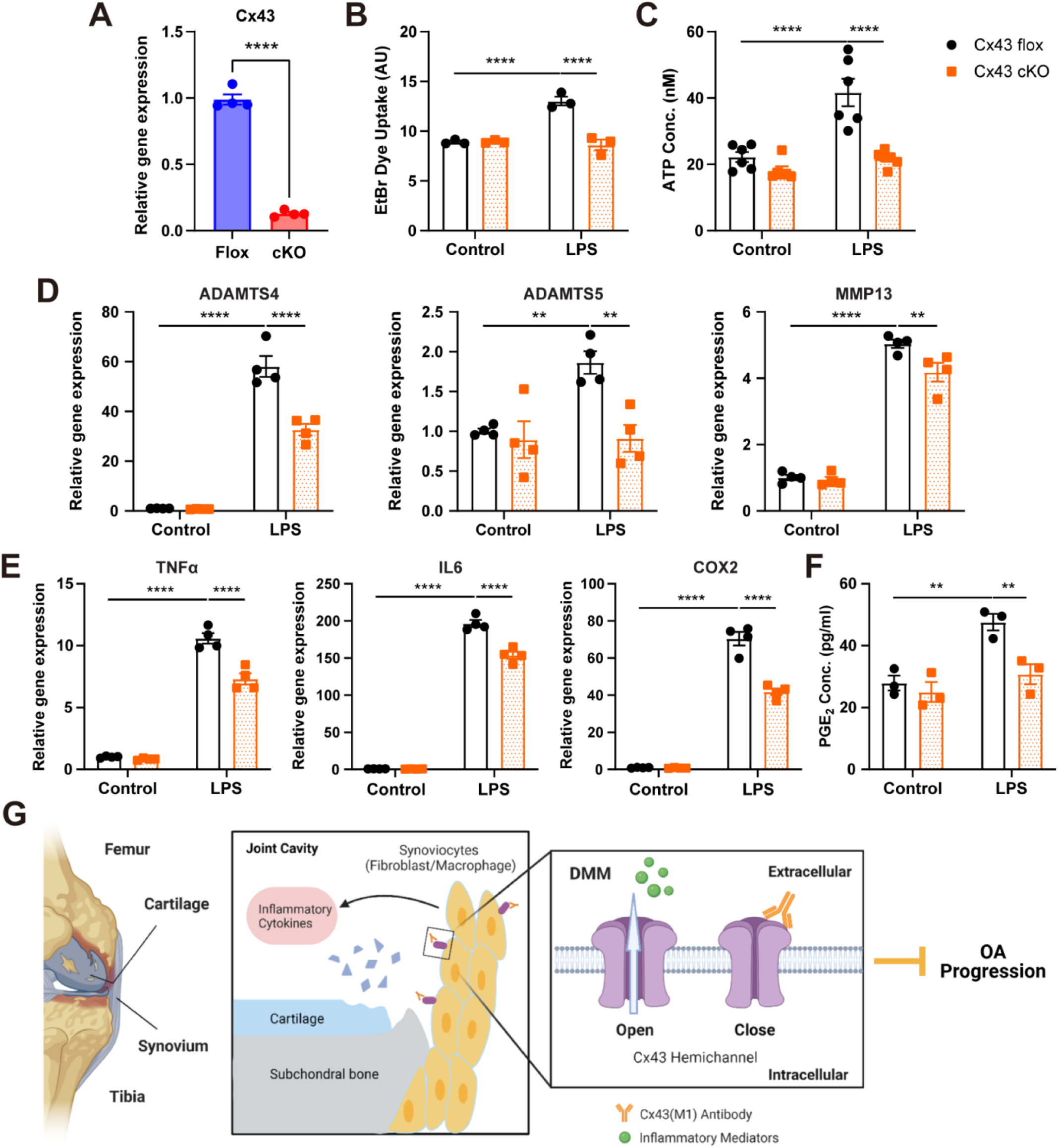
Cx43 deletion in bone marrow-derived macrophages (BMDM) reduces LPS-induced inflammatory gene expression, accompanied by decreased ATP and PGE₂ release. **(A)** BMDMs were isolated from Cx43 flox/flox or Cx43 cKO mice and analyzed for Cx43 expression. n = 4 mice/group. **(B)** BMDMs from Cx43 flox/flox or Cx43 cKO mice were treated with LPS, and EtBr dye uptake was quantified by fluorescence microscopy and NIH ImageJ. **(C)** Extracellular ATP release was determined in LPS-treated BMDMs. **(D,E)** RNA was extracted and subjected to qRT-PCR analysis for ADAMTS4, ADAMTS5, MMP13, TNFα, IL6, and COX2 mRNA expression. **(F)** Extracellular PGE_2_ release was determined in LPS-treated BMDMs. n =3-6/group. Data are presented as mean ± SEM. **, p < 0.01; ****, p < 0.0001. **(G)** Proposed model: During OA progression, activated synovial cells in the inflamed synovium produce catabolic and pro-inflammatory mediators that promote proteolytic enzyme production amd drive cartilage breakdown. The synovium of knee joints comprises macrophages and fibroblasts that express Cx43. Administration of the Cx43 antibody blocks hemichannel opening and reduces the release of inflammatory mediators, thereby mitigating cartilage breakdown, subchondral bone sclerosis, and pain-related behaviors.

To further validate these findings in an independent macrophage population, we also examined primary peritoneal macrophages isolated from Cx43 flox/flox and cKO mice. Consistent with the results observed in BMDMs, Cx43 expression was reduced by approximately 90% in Cx43 cKO macrophages (Fig. S7A), and Cx43 deletion significantly decreased LPS-induced extracellular ATP release (Fig. S7B). Moreover, Cx43 deficiency suppressed the expression of inflammatory and catabolic genes, including MMP13, TNFα, IL6, and COX2 (Fig. S7C, D), along with reduced PGE₂ production (Fig. S7E). Collectively, these data demonstrate that Cx43 regulates ATP- and PGE_2_-dependent inflammatory signaling across multiple macrophage populations, providing a mechanistic basis for the protective effects of Cx43 deletion or inhibition in OA.

Based on these findings, we propose a model in which synovial Cx43 HCs act as key mediators of joint inflammation and structural deterioration in OA, highlighting the crucial role of Cx43 HCs in regulating pro-inflammatory mediators under OA conditions. Selective targeting of synovial Cx43 HCs with a monoclonal antibody offers a promising strategy to mitigate OA progression by attenuating inflammatory signaling, preserving joint structure, and alleviating pain (Fig. 7G). Together, these findings position synovial macrophage Cx43 HCs as a mechanistically defined therapeutic target and support a shift toward inflammation-directed intervention in OA.

## Materials and Methods

### Animals and surgery models

C57BL/6J and LysM-cre mice were obtained from The Jackson Laboratory (stock No. 000664 and 004781, respectively). Mice with a floxed Cx43 gene (Cx43 flox/flox) were originally generated by Dr. Klaus Willecke (University of Bonn, Germany) [30] and were provided by Dr. Roberto Civitelli (Washington University, USA). Macrophage-specific Cx43 conditional knockout mice were generated by interbreeding Cx43 flox/flox mice with mice carrying the LysM-cre transgene. Littermate controls were used for all experiments. Male mice at 12 weeks of age were used unless otherwise specified. All mice were maintained on a C57BL/6J background. Mice were housed under specific pathogen-free conditions in a temperature-controlled facility with a 12-hour light/dark cycle at the UTHSCSA Laboratory Animal Resources (LAR) facility, with *ad libitum* access to food and water. All animal procedures were approved by the UTHSCSA Institutional Animal Care and Use Committee (IACUC).

DMM surgery was performed on the right knee joint of mice as previously described [31]. Briefly, a medial parapatellar incision was made in the right knee joint to expose the joint capsule. The medial meniscotibial ligament was identified and transected to destabilize the medial meniscus. Sham-operated mice underwent the same surgical procedure without ligament transection. Mice were randomly assigned to experimental groups.

### AAV vector and intra-articular injection

Recombinant AAV5 vectors were used for *in vivo* gene delivery based on their high transduction efficiency in joint tissues [32, 33]. AAV5-GFP (control) and AAV5-GFP-iCre were purchased from Vector Biolabs (Malvern, PA, USA). The iCre construct encodes an improved Cre recombinase co-expressed with enhanced green fluorescent protein (eGFP) under the control of a CMV promoter, with Cre and eGFP separated by a 2A self-cleaving peptide.

For intra-articular administration, mice were anesthetized and a longitudinal skin incision was made to expose the patellar ligament and patella. A total of 5 × 10⁹ viral particles in a 10 μL volume were injected into the knee joint cavity by inserting a fine needle beneath the patella of the right knee. AAV vectors were administered 7 days prior to DMM or sham surgery to allow sufficient transgene expression. Successful transduction and distribution of AAV expression in joint tissues were confirmed by GFP fluorescence.

### Cx43(M1) antibody generation and treatment

A monoclonal Cx43(M1) antibody targeting the second extracellular loop of Cx43 was originally generated by Abmart (Tulsa, OK, USA) and characterized as previously described [20]. Briefly, mice were immunized with a peptide corresponding to the extracellular domain of Cx43. Following hybridoma screening and functional characterization, genes encoding the antibody heavy- and light-chain variable regions were amplified from the hybridoma cell line by RT-PCR and cloned into expression vectors. The constructs were co-transfected into HEK293F cells, and antibodies were purified from culture supernatants using protein A affinity chromatography. For *in vivo* studies, Cx43(M1) antibody was administered by intraperitoneal injection at a dose of 25 mg/kg. For the standard treatment regimen, mice received Cx43(M1) or vehicle (phosphate-buffered saline, PBS) 30 minutes and 4 weeks following sham or DMM surgery. To evaluate the effects of a single administration, mice received a one-time intraperitoneal injection of Cx43(M1) (25 mg/kg) or PBS 30 minutes after surgery. For delayed-treatment experiments, the initial dose of Cx43(M1) (25 mg/kg) was administered 10 days after DMM surgery, followed by additional injections on days 25 and 40. Mice were sacrificed at 8 weeks post-surgery for tissue analysis.

### Histology

At 8 weeks following induction of OA, joint samples from DMM and sham-operated mice were harvested, fixed in 4% paraformaldehyde for 48 hours, and decalcified in 10% EDTA (pH 7.5) for 3 weeks. Tissues were then embedded in paraffin, and 5-μm-thick sections were prepared. For histological evaluation, sections were deparaffinized in xylene and rehydrated through a graded ethanol series. Cartilage integrity was assessed by Safranin O/Fast Green staining, while general joint morphology and synovial structure were evaluated using hematoxylin and eosin (H&E) staining. Cartilage degeneration was quantified using the Osteoarthritis Research Society International (OARSI) scoring system as previously described [34]. Both the medial femoral condyle (MFC) and medial tibial plateau (MTP) were evaluated using three representative sections per joint, and OA severity was expressed as the average score. Synovial inflammation was assessed based on synovial membrane thickening and hyperplasia and was graded on a scale of 0-3 (0, no synovitis; 1, mild; 2, moderate; 3, severe) as previously described [35, 36]. All histological evaluations were performed in a blinded manner.

### Immunohistochemistry and immunofluorescence staining

For immunohistochemistry staining, paraffin-embedded joint sections (5 μm) were deparaffinized, rehydrated, and subjected to antigen retrieval. Sections were then incubated overnight at 4 °C with primary antibodies against MMP13 (ab39012, Abcam, Cambridge, UK) or collagen X (14-9771-82, Thermo Fisher Scientific, Waltham, MA, USA). After washing, sections were incubated with biotinylated secondary antibodies followed by avidin-biotin-peroxidase complex (ABC) reagent. Signals were visualized using DAB substrate (SK4100, Vector Laboratories, Newark, CA, USA), and nuclei were counterstained with hematoxylin. Images were captured using an optical microscope (BZ-X710, Keyence, Japan). The percentage of MMP13- or collagen X-positive chondrocytes was quantified using ImageJ software.

For immunofluorescence staining, joint tissues were fixed in 4% paraformaldehyde, decalcified in 10% EDTA, cryoprotected in 30% sucrose, and embedded in Tissue-Tek CRYO OCT compound (14-373-65, Andwin Scientific, Simi Valley, CA, USA). Cryosections (12 μm) were permeabilized and blocked, followed by overnight incubation at 4 °C with primary antibodies against Cx43 (CT) (1:100) [37], GFP (ab290, Abcam, Cambridge, UK), CD68 (MCA1957, Bio-Rad Laboratories, Hercules, CA, USA), iNOS (PA1036, Thermo Fisher Scientific, Waltham, MA, USA), and Arg1 (NB10059740, Novus Biologicals, Littleton, CO, USA). Sections were then incubated with appropriate fluorescent secondary antibodies. For antibody localization, Alexa Fluor-conjugated anti-human IgG secondary antibody was used to detect the distribution of administered Cx43 antibody. Slides were mounted with antifade mounting medium and imaged using a confocal microscope (LSM770, Carl Zeiss, Germany).

### Micro-computed tomography

Subchondral bone structure was evaluated using 3D reconstruction from a μCT imaging system (Brüker SkyScan 1173; Brüker microCT, Kontich, Belgium). Samples were scanned in saline with the following parameters: 60 kV, 167 μA beam intensity, 0.5 mm aluminum filter, 0.7° rotation step, 4-frame averaging, 1090 ms integration time, 1024 × 1024-pixel matrix, and a 10 μm isotropic voxel size. After scanning, noise was removed from the images by eliminating disconnected objects smaller than 4 pixels in size. Morphometric analysis was performed on 50 slices extending proximally, beginning with the first slice in which the tibial condyles had fully merged. The subchondral bone was manually segmented from the cortical shell on key slices using a contouring tool, and the contours were automatically morphed to segment the trabecular bone on all slices. The morphometry was reconstructed and analyzed. A grayscale value of 95 in a set of 8-bit slices was set as the threshold and was applied to all specimens after comparing grayscales and binarized images in both groups. After thresholding, the BV/TV (%), Tb.Th (mm), and Tb.N (mm^−1^) were quantified.

### Animal behavioral tests

The paw withdrawal threshold was measured using von Frey filaments ranging from 0.007 g to 2 g. Mice were placed in acrylic chambers (5.5 × 10 cm) suspended above a wire mesh grid and allowed to acclimate for 30 min before testing. Von Frey filaments were applied perpendicularly to the plantar surface of the hind paw until the filament bent, and held in place for up to 3 s. A positive response was recorded if the paw was sharply withdrawn or if the mouse flinched immediately upon filament removal, as previously described [38, 39]. Testing began with the 0.4 g filament and progressed according to the up-down method.

Spontaneous locomotor activity was analyzed using the open-field test. Mice were placed into the center of a chamber (25 × 25 × 25 cm) to allow explore freely. The experiments were performed for 10 minutes. Total distance traveled and immobility time were measured using ANY-maze behavior tracking software (Stoelting Co., Wood Dale, IL, USA).

### *In vivo* dye uptake assay

We developed an approach to assess HC activity *in vivo* [40]. Briefly, Evans blue dye (MW: 900 Da) (200 mg/kg) and FITC-dextran (MW: 10,000 Da) (200 mg/kg) were co-injected via the tail vein. One-hour post administration of the dye, mice were anesthetized, and cardiac perfusion was performed with cold PBS and 4% paraformaldehyde. Both hindlimbs were isolated and prepared for frozen sectioning. Sections were stained with DAPI and photographed using a Keyence microscope (BZ-X710, Keyence, Osaka, Japan). The intensity of dye uptake was quantified using NIH Image J software.

### Isolation of primary human synovial macrophages

Primary human synovial macrophages were isolated from synovial tissue obtained during knee joint replacement surgery. This study was reviewed and determined to be exempt by the Institutional Review Board (IRB). Cartilage and synovial tissue samples used in this study were surgical discards, and no informed consent was required in accordance with the approved IRB protocol, as no identifiable patient information was collected. Only limited de-identified information, including age, sex, and clinical diagnosis (e.g., osteoarthritis), was recorded. Fresh synovial tissues were collected in cold PBS containing antibiotics and processed under sterile conditions. After removal of adipose tissue and debris, samples were minced into small fragments (1-2 mm) and digested in collagenase IV solution (2 mg/mL in complete DMEM/F12) at 37 °C with gentle agitation for 2-4 hours. The resulting cell suspension was filtered through a 70-μm cell strainer and centrifuged to obtain a single-cell suspension. Cells were washed with PBS containing 0.5% BSA and resuspended. Synovial macrophages were purified using magnetic-activated cell sorting (MACS) with anti-CD14-conjugated magnetic beads according to the manufacturer’s instructions. The purified macrophages were collected, and cell purity was validated prior to downstream analyses.

### Isolation of peritoneal macrophages and bone marrow-derived macrophages

Peritoneal macrophages were elicited by intraperitoneal injection of 2 mL of 4% thioglycollate medium (BD Biosciences, San Jose, CA, USA) into Cx43 flox/flox and cKO mice. Four days after injection, cells were harvested by peritoneal lavage using 5 mL of DMEM/F12 medium. Recovered cells were cultured at 37 °C in a humidified incubator with 5% CO₂ in DMEM/F12 supplemented with 10% fetal bovine serum and 1% penicillin/streptomycin.

Bone marrow-derived macrophages were isolated from the femurs and tibiae of Cx43 flox/flox and cKO mice. Bone marrow cells were flushed from the bones and cultured in DMEM/F12 medium containing 10% FBS, 1% penicillin/streptomycin, and 15% conditioned medium from L929 mouse fibroblasts as a source of macrophage colony-stimulating factor (M-CSF). Cells were differentiated into macrophages over 7 days prior to use.

### Cell culture and inflammatory stimulation

RAW264.7 cells and SW982 cells were maintained in Dulbecco’s Modified Eagle Medium (DMEM). All culture media were supplemented with 10% fetal bovine serum (FBS) and 1% penicillin/streptomycin. Cells were maintained at 37 °C in a humidified incubator with 5% CO₂. For inflammatory stimulation experiments, RAW264.7 macrophages were treated with 100 ng/ml lipopolysaccharide (LPS) for 4 hours to induce inflammatory responses, while SW982 synovial fibroblasts were stimulated with 1 ng/ml interleukin-1β (IL-1β) for 14 hours. Where indicated, cells were pretreated with 2 μg/ml Cx43(E2) prior to stimulation. After treatment, cells and conditioned media were collected for downstream analyses.

### Extracellular PGE_2_ and ATP Measurements

The conditioned media was collected and centrifuged to remove cell debris. The level of extracellular PGE_2_ was determined using a PGE_2_ ELISA kit according to the manufacturer’s protocol (514010, Cayman Chemical, Ann Arbor, MI, USA). To assess ATP release, 100 μM ARL67156 (an ecto-ATPase inhibitor) was added to the media to prevent ATP hydrolysis. Extracellular ATP concentrations were then measured using a luciferin/luciferase-based ATP Determination Kit (#A22066, Thermo Fisher Scientific, Waltham, MA, USA) according to the manufacturer’s protocol.

### *In vitro* dye uptake assay

Cells were incubated with 0.1 mM ethidium bromide (EtBr; MW 394 Da) and 1 mg/mL FITC-dextran (MW 10 kDa) for 5 minutes at room temperature. EtBr was used as a permeable tracer to assess HC activity, whereas FITC-dextran, which is excluded from intact cells due to its large molecular size, served as a negative control for nonspecific uptake associated with cell membrane damage. Following incubation, cells were washed five times with PBS and fixed with 2% paraformaldehyde for 10 minutes. Fluorescence images were acquired using a 20× objective on a Keyence BZ-X710 fluorescence microscope (Keyence, Osaka, Japan). For each experimental condition, at least six random fields were captured, and a minimum of 30 cells per image was analyzed. EtBr fluorescence intensity was quantified using ImageJ software (NIH, Bethesda, MD, USA), and results were expressed as mean fluorescence intensity (arbitrary units). All experiments were performed in triplicate.

### RNA isolation and real-time PCR

Total RNA was isolated from RAW264.7 and SW982 cells using TRI Reagent (Molecular Research Center, Cincinnati, OH, USA) according to the manufacturer’s instructions. RNA concentration and purity were determined using a NanoDrop 2000 spectrophotometer (Thermo Fisher Scientific, Waltham, MA, USA). cDNA was synthesized using the High-Capacity RNA-to-cDNA Kit (43-889-50, Applied Biosystems, Bedford, MA, USA). Real-time PCR was performed using an ABI 7900 PCR system and SYBR Green (1725124, Bio-Rad Laboratories, Hercules, CA, USA) with a two-step protocol (94°C for 15 seconds and 60°C for 60 seconds). The ΔΔC_T_ method was used for qPCR data analysis. GAPDH was used as housekeeping gene control. Experiments were run in triplicate. The primer sequences used in this study are listed in Table S1.

### Statistical analysis

Statistical analysis was performed using GraphPad Prism 8 statistics software (GraphPad Software, San Diego, CA, USA). All data are presented as the mean ± SEM. Student’s t-test, one-way ANOVA and two-way ANOVA analysis of variance with Tukey’s test and multiple comparisons were used for statistical analysis. Asterisks indicate the degree of significant differences compared with the controls (*P < 0.05; **P < 0.01; ***P < 0.001; ****P < 0.0001).

## Discussion

In this study, we identify Cx43 HCs as critical regulators of synovial inflammation and joint degeneration in OA. Using complementary genetic and pharmacological approaches, we demonstrate that inhibition of Cx43 HC activity attenuates cartilage degradation, synovial inflammation, subchondral bone remodeling, and pain-associated behaviors in a murine OA model. Importantly, our findings identify synovial macrophages as a key cellular target of Cx43-mediated signaling and establish a mechanistic link between Cx43 HC activity, ATP- and PGE₂-dependent inflammatory responses, and OA progression.

Previous studies have linked elevated Cx43 expression in OA to chondrocyte dysfunction, including chondrocyte senescence, chondrocyte-mesenchymal transition, and the propagation of degenerative signals through Cx43-enriched extracellular vesicles [41, 42]. Cx43 silencing has also been shown to ameliorate disease severity in inflammatory arthritis models [43]. However, these studies have largely focused on chondrocyte-associated Cx43 or total Cx43 expression. Our study addresses this critical gap by identifying synovial macrophage Cx43 HCs as functional drivers of OA progression. By selectively blocking Cx43 HCs without disrupting gap junctional communication, and by using macrophage-specific Cx43 deletion, we show that macrophage Cx43 HCs regulate synovial inflammation, cartilage degeneration, subchondral bone sclerosis, and pain-related behaviors. This cell- and channel-specific mechanism shifts the focus of Cx43 biology in OA from chondrocyte-centered degeneration toward macrophage-mediated inflammatory joint pathology.

The use of complementary genetic and pharmacological approaches allowed us to distinguish the pathological contribution of Cx43 HCs from broader effects of total Cx43 expression. Cx43(M1) targets the second extracellular loop of Cx43, a region highly conserved across mouse, rat, and human, and has been shown in previous studies to selectively block HCs while preserving physiological gap junction function in astrocytes and osteocytes [20, 21]. Cx43(M1)-mediated HC-inhibition has been associated with improved recovery from spinal cord injury and ocular nerve injury [20, 44]. In the present OA model, selective inhibition of HCs was sufficient to protect against cartilage degeneration, synovial inflammation, subchondral bone sclerosis, and pain-related behaviors, supporting aberrant HC opening as a key contributor to inflammatory joint degeneration. Importantly, neither pharmacological inhibition of Cx43 HCs nor genetic deletion of Cx43 resulted in detectable baseline functional deficits in sham-operated mice, suggesting pathological HC activity can be targeted without broadly disrupting homeostatic joint function. The therapeutic relevance of this approach is further supported by the efficacy of both early and delayed Cx43(M1) administration. Delayed treatment attenuated OA progression even after cartilage degeneration had begun, indicating a therapeutically relevant intervention window. In addition, a single injection of Cx43(M1) produced sustained protective effects and antibody signal remained detectable in the synovium for at least two weeks. This prolonged synovial retention may improve therapeutic practicality by reducing dosing frequency while maintaining target engagement.

The delivery profile of biologic therapeutics is a critical consideration in OA, particularly because specific targeting of chondrocytes remains challenging. The dense extracellular matrix and avascular nature of articular cartilage limit the penetration of large molecules [26]. Our *in vivo* binding studies showed that systemically administered Cx43 monoclonal antibody predominantly accumulated in the synovium, with minimal distribution in cartilage. Thus, rather than requiring direct chondrocyte targeting, Cx43(M1) appears to act primarily through modulation of the synovial compartment. By blocking Cx43 HCs on inflammatory synovial cells, the antibody may interrupt upstream inflammatory signing before synovial-derived mediators amplify cartilage matrix degradation and subchondral bone remodeling. This synovium-centered mechanism is consistent with growing evidence that synovial inflammation can occur before cartilage degeneration with infiltration of mononuclear cells, thickening of the synovial lining layer, and production of inflammatory cytokines [45]. Our findings further support the concept that selective targeting of the synovial compartment may be sufficient to modify disease progression despite limited cartilage penetration. In agreement with this view, Gui et al. [46] reported that large nanoparticles (∼120 nm) encapsulating superoxide dismutase predominantly accumulated in the synovium and failed to penetrate into the articular cartilage. Despite the lack of direct cartilage contact, these synovium-targeted nanoparticles effectively attenuated cartilage degradation, reduced catabolic protease expression including Mmp13 and Adamts5, and mitigated synovitis in the DMM model. Therefore, synovium represents a highly accessible and therapeutically relevant compartment for biologic interventions in OA and supports a model in which modulating the synovial inflammation exerts protective effects across the entire joint.

Our findings position synovial macrophages as a key cellular site of Cx43 HC activity in OA. Cx43 HC-blockade reduced iNOS⁺/CD68⁺ macrophage accumulation *in vivo*, inhibited LPS-induced HC opening, inflammatory gene expression, and ATP and PGE₂ release in both primary human OA synovial macrophages and a mouse macrophage cell line. Moreover, macrophage-specific Cx43 deletion recapitulated the protective effects of antibody treatment. These convergent findings indicate that macrophage Cx43 HCs are not only associated with synovial inflammation but functionally contribute to inflammatory joint degeneration. This conclusion is consistent with the established pathogenic role of macrophages in OA. Activated macrophages are present in the majority (∼ 76%) of OA knees, and their abundance correlates with key symptoms including knee pain, joint space narrowing, and osteophytes [41]. Functional evidence further supports their pathogenic contribution: depletion of macrophages from synovial cell cultures using anti-CD14-conjugated magnetic beads reduced the production of IL-1, TNF-α, MMPs, and aggrecanase-all of which mediate destructive processes in OA [5]. Likewise, genetic disruption of macrophage-associated inflammatory pathways has been shown to attenuate OA-associated bone remodeling, as CD14 knockout mice are protected from both age-related and post-surgical changes in subchondral BMD and trabecular thickness [42]. Increased M1 polarization of synovial macrophages has also been shown to exacerbate experimental OA, in part through the secretion of Rspo2 and activation of β-catenin signaling in chondrocytes [43]. Together, these observations place macrophages at the center of OA-associated inflammatory crosstalk and support our conclusion that macrophage Cx43 HCs act as upstream regulators of synovial inflammation and joint degeneration.

Mechanistically, our findings support a model in which Cx43 HCs regulate inflammatory signaling through the release of ATP, thereby amplifying inflammatory cascades within the synovium. In this context, Cx43 HCs may act as upstream gatekeepers of ATP availability, controlling the magnitude and duration of downstream purinergic signaling pathways in OA joints. Extracellular ATP functions as a key danger-associated molecular pattern (DAMP) that activates purinergic receptors, particularly P2X7, leading to inflammasome activation and production of pro-inflammatory cytokines such as IL-1β [47, 48]. ATP signaling is also closely linked to macrophage polarization [49, 50]. Elevated extracellular ATP promotes a pro-inflammatory (M1-like) phenotype, characterized by increased expression of iNOS and secretion of inflammatory mediators, while also enhancing NLRP3 inflammasome activation. In turn, activated M1 macrophages release additional ATP through channels such as connexin HCs and pannexin-1, potentially creating a feed-forward loop that sustains macrophage activation and propagates synovial inflammation. Our findings support this model and suggest that Cx43 HC-dependent ATP release may serve as an upstream amplifier of macrophage-driven inflammatory signaling in OA.

However, recent work highlights a more complex and context-dependent role for PGE₂ in joint biology. A recent study demonstrated that the prostaglandin-degrading enzyme 15-hydroxyprostaglandin dehydrogenase (15-PGDH) is upregulated in aged and OA cartilage, where it reduces endogenous PGE₂ levels and is associated with cartilage degeneration [51]. Notably, pharmacological inhibition of 15-PGDH increased PGE₂ levels, promoted cartilage regeneration, and reduced inflammatory cytokine production. These findings suggest that physiological levels of PGE₂ may exert protective or regenerative effects, whereas excessive, sustained production may contribute to inflammation and tissue damage. In this context, our findings suggest that Cx43 HCs regulate PGE₂ release from activated synovial macrophages, linking HC activity to inflammatory amplification and pain sensitization within the synovial compartment. Thus, the pathological impact of Cx43 HC activity may depend not only on the amount of PGE₂ released but also on its cellular source, tissue localization, and balance with degradation pathways such as 15-PGDH. Importantly, the emerging complexity of PGE₂ biology suggests that therapeutic approaches may need to consider not only suppression of inflammatory mediators but also preservation of their homeostatic functions.

Several limitations of this study should be acknowledged. While our data strongly support a role for ATP and PGE₂ as downstream mediators, the precise intracellular signaling pathways linking Cx43 HC activation to inflammatory gene expression remain to be fully defined. Future studies examining downstream purinergic signaling, inflammasome activation, and transcriptional regulators may further clarify how Cx43 HC activity sustains inflammatory responses within the synovium. In addition, although macrophages appear to be the primary drivers, contributions from other joint-resident cells cannot be excluded. Finally, the DMM model primarily reflects post-traumatic OA driven by mechanical instability. Whether Cx43 HC inhibition is similarly effective in age-associated, metabolic or obesity-related OA models remain an important question for future investigation.

In conclusion, this study establishes Cx43 HCs as key mediators of synovial inflammation and joint degeneration in OA. By demonstrating that selective inhibition of Cx43 HCs attenuates cartilage degradation, synovitis, subchondral bone pathology, and pain-related behaviors, our findings identify macrophage-associated Cx43 HC signaling as a mechanistically defined contributor to OA progression. The efficacy of delayed antibody administration, its preferential localization to synovial macrophages, and the recapitulation of protective effects in macrophage-specific Cx43 knockout mice all support the clinical translatability of this approach. More broadly, these findings shift the focus from a predominantly chondrocyte-centered view of OA toward synovial macrophage Cx43 HCs as mechanistically defined and therapeutic targets. Future studies should focus on optimizing delivery routes, evaluating long-term safety, and testing efficacy in additional preclinical models that reflect the heterogeneity of human OA.

## Acknowledgement

We thank Di Chen for valuable suggestions and insightful discussions throughout this study. We also thank Klaus Willecke for generously providing the Cx43 flox mice. This work was supported by the Welch Foundation (Grant No. AQ-1507).

## Author contributions

J.X.J. conceived and supervised the study. R.H. and J.X.J. designed the study. R.H., Y.T., X.W., L.M., M.A.R., H.X., L.Z., T.G., J.Z., and S.G. performed experiments and/or analyzed data. F.A.B. and A.A. provided human tissue samples from OA patients. R.H. and J.X.J. wrote the manuscript. All authors read, reviewed, and approved the final manuscript.

