## Supplemental Materials for "Targeting Macrophage Connexin 43 Hemichannels Suppresses Synovial Inflammation and Osteoarthritis"

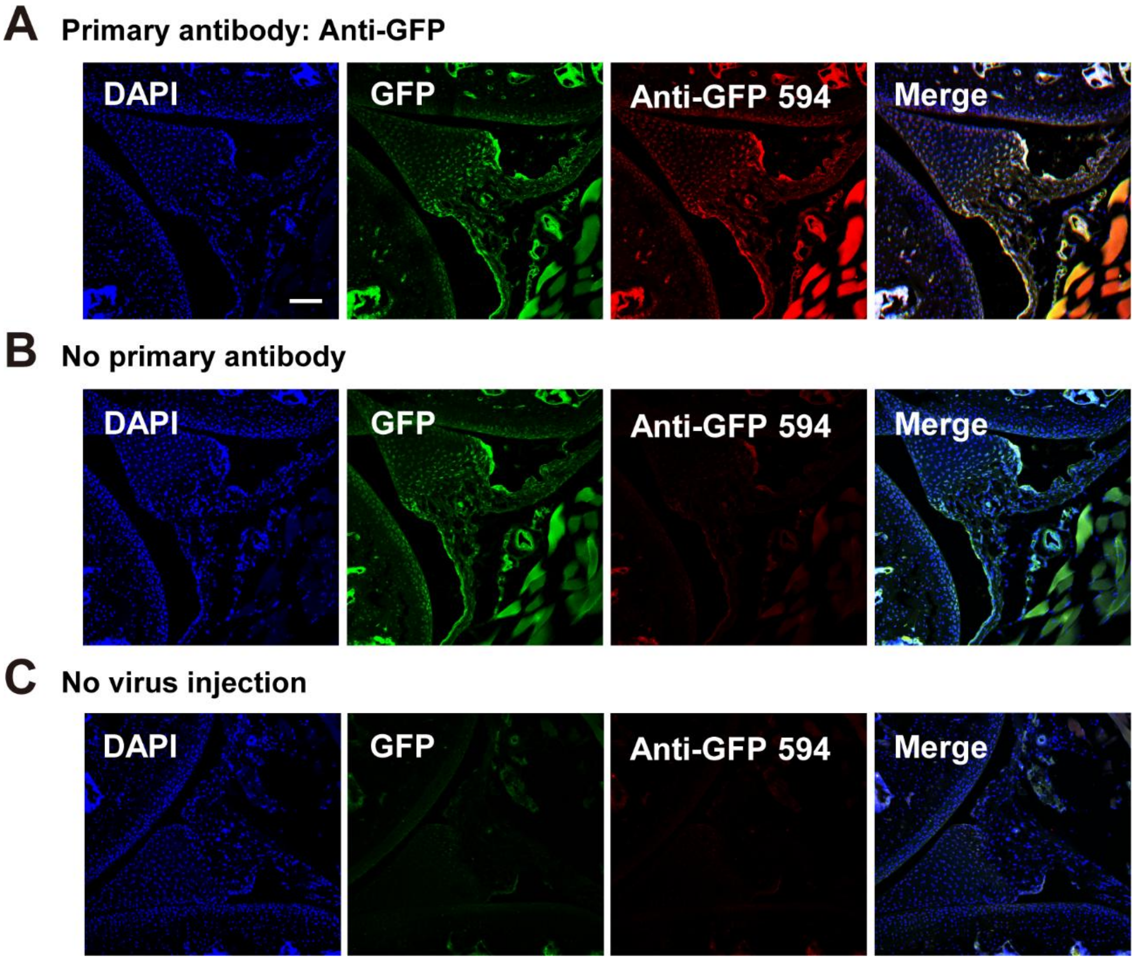


**Figure S1. Immunofluorescent detection of GFP expression in joint tissues after intra-articular AAV-GFP administration.**

**(A)** Immunofluorescence staining revealed GFP expression in the synovium and articular cartilage of knee joints of 4-month-old C57BL/6 mice that received intra-articular AAV5-GFP injections at 2 months of age. Expression was visualized by direct GFP fluorescence (green channel) or by anti-GFP immunostaining (red channel). Scale bar, 50 µm. **(B)** No-primary antibody control showing GFP signal without anti-GFP immunolabeling. Background-level signal is observed in the red channel. **(C)** No virus injection negative control showing background-level signal in both green and red channels.

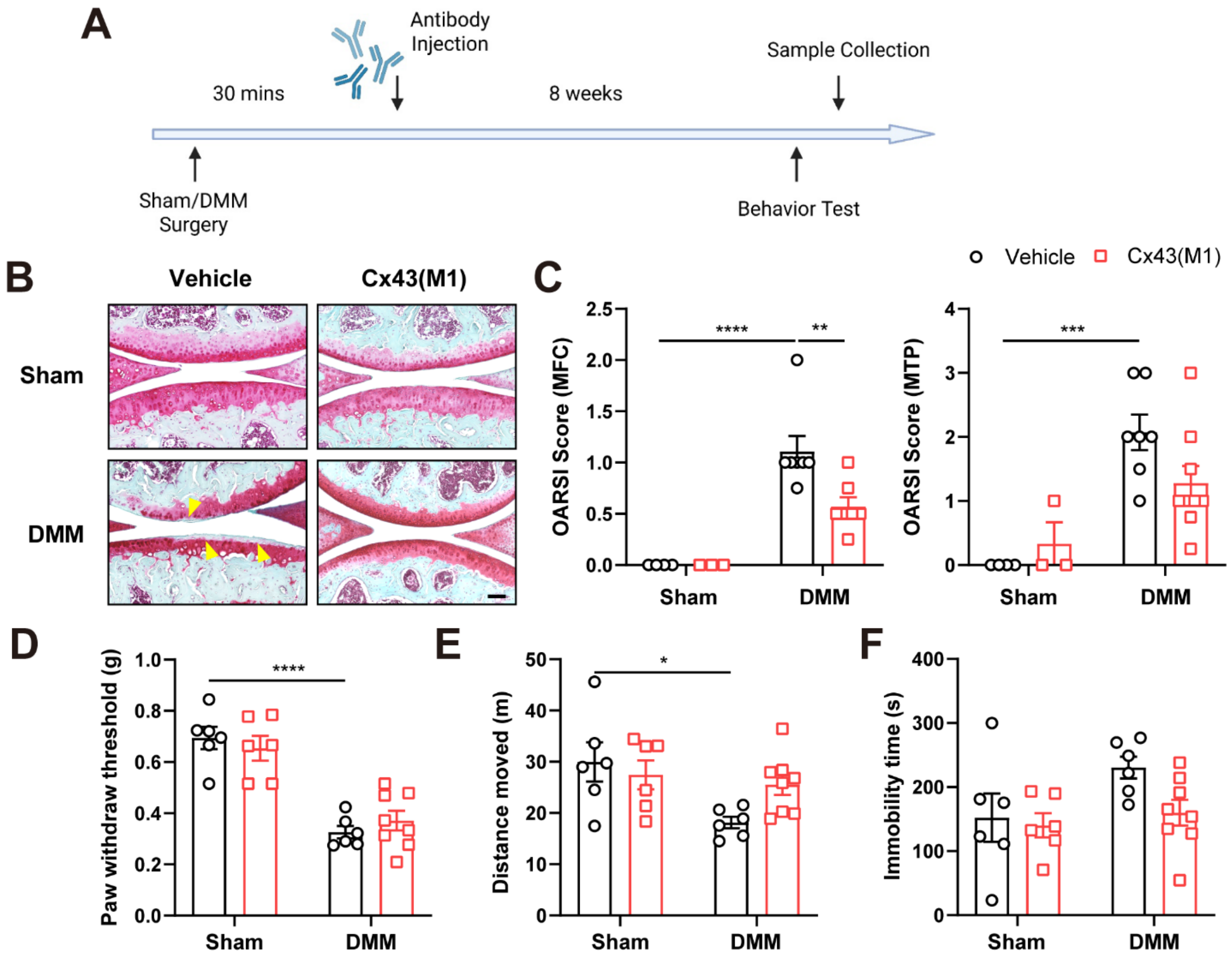


**Figure S2. Single-dose Cx43 antibody treatment preserved cartilage integrity and reduced OA-associated pain.**

**(A)** Schematic diagram of the timeline for surgical procedures, antibody injections, and outcome assessments. The Cx43 hemichannel-blocking antibody was administered 30 minutes post-surgery. Behavioral tests were conducted, and tissue samples were collected 8 weeks after surgery. **(B)** Representative images of Safranin O/Fast Green staining in vehicle- or Cx43(M1) antibody-treated mice after sham or DMM surgery. Areas of cartilage destruction are indicated by yellow arrowheads. Scale bar, 200 µm. **(C)** Averaged OARSI scores of the medial femoral condyle (MFC, left panel) and medial tibial plateau (MTP, right panel). n = 3-7 mice/group. **(D)** Hind paw withdrawal thresholds were measured using the von Frey filament test at 8 weeks post-surgery. **(E-F)** Open field behavioral assessments were conducted at 8 weeks post-surgery. n = 6-8 mice/group. Data are presented as mean ± SEM. *, $p < 0.05$; **, $p < 0.01$; ***, $p < 0.001$; ****, $p < 0.0001$.

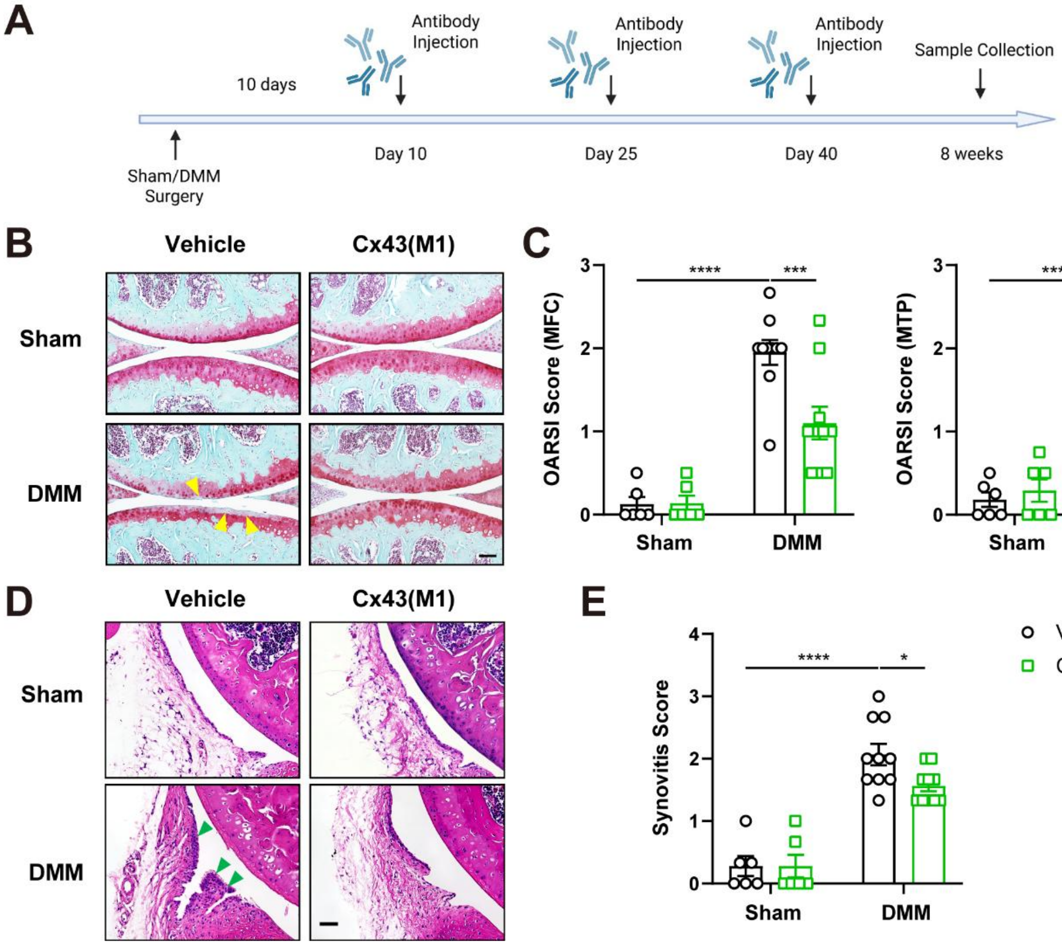


**Figure S3. Delayed Cx43 antibody treatment attenuated articular cartilage breakdown and decreased synovium inflammation.**

**(A)** Schematic diagram of the timeline for surgical procedures, antibody injections, and outcome assessments. The Cx43 hemichannel-blocking antibody was administered at 10, 25, and 40 days post-surgery. Tissue samples were collected 8 weeks after surgery. **(B)** Representative images of Safranin O/Fast Green staining in vehicle- or Cx43(M1) antibody-treated mice after sham or DMM surgery. Areas of cartilage destruction are indicated by yellow arrowheads. Scale bar, 200 μm. **(C)** Averaged OARSI scores of the medial femoral condyle (MFC, left panel) and medial tibial plateau (MTP, right panel). n = 6-10 mice/group. **(D-E)** Representative images of H&E staining **(D)** and quantification of synovitis scores **(E)** in vehicle- or Cx43(M1) antibody-treated mice after sham or DMM surgery. Areas of synovial inflammation are indicated by green arrowheads. Scale bar, 200 μm. n = 6-11 mice/group. Data are presented as mean ± SEM. *, $p < 0.05$; ***, $p < 0.001$; ****, $p < 0.0001$.

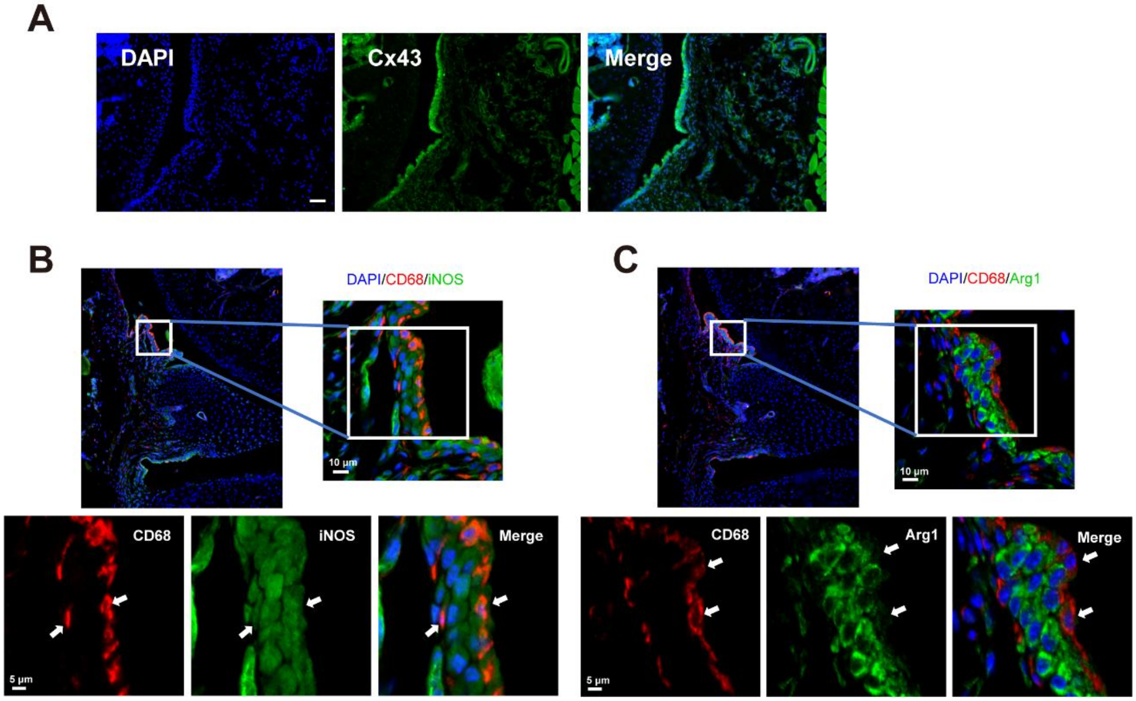


**Figure S4. Pro-inflammatory M1-type macrophages are the predominant macrophage population increased in DMM-induced OA.**

**(A)** Tissue sections were immunolabeled with Cx43 antibody. Scale bar, 50 µm. **(B)** Pro-inflammatory M1-like macrophages were identified by co-immunofluorescence staining for CD68 (red channel) and iNOS (green channel), with one synovial region shown at higher magnification in the lower panels. White arrows indicate $CD68^+$/$iNOS^+$ double-positive cells. **(C)** Tissue sections were co-immunolabeled with CD68 (red channel) and the anti-inflammatory M2-like macrophages marker Arg1(green channel). White arrows indicate $CD68^+$ cells that are negative for Arg1 expression.

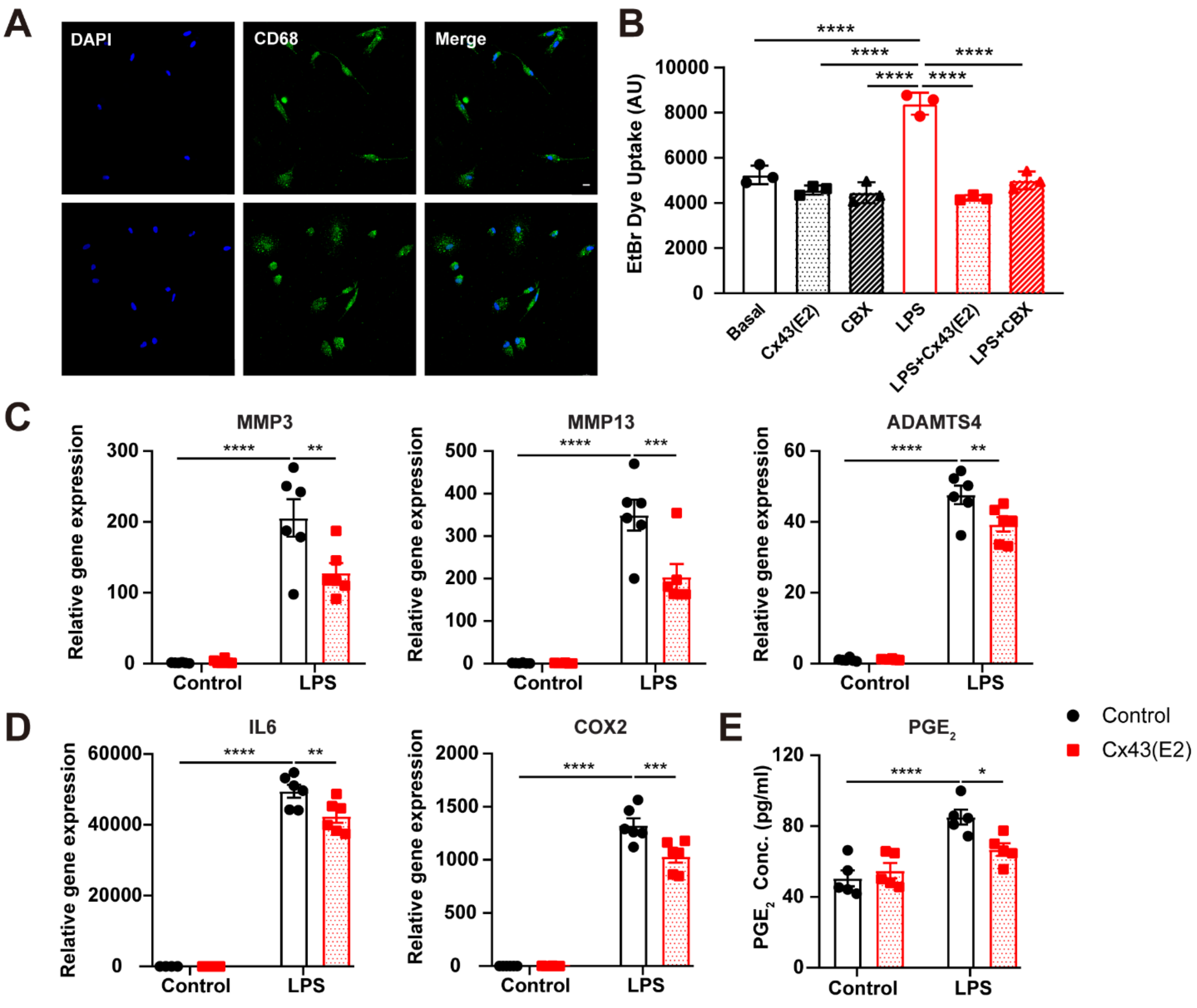


**Figure S5. Cx43 hemichannel-blocking antibody inhibited hemichannel opening and suppressed inflammatory responses in mouse macrophage cell line R264.7.**

**(A)** Purified human OA synovial macrophages were immunolabeled with CD68 antibody. Scale bar, 10 μm. **(B)** Mouse macrophage R264.7 cells were treated with LPS and Cx43(E2) antibody or CBX. EtBr dye uptake was quantified by fluorescence microcopy and NIH ImageJ. **(C-D)** RNA was extracted from R264.7 cells treated with LPS and Cx43(E2) antibody and subjected to qRT-PCR analysis for MMP3, MMP13, ADAMTS4, IL6, and COX2 mRNA expression. **(E)** Extracellular $PGE_2$ release was measured in mouse macrophages treated with LPS and Cx43(E2) antibody. n = 3-6/group. Data are presented as mean ± SEM. *, $p < 0.05$; **, $p < 0.01$; ***, $p < 0.001$; ****, $p < 0.0001$.

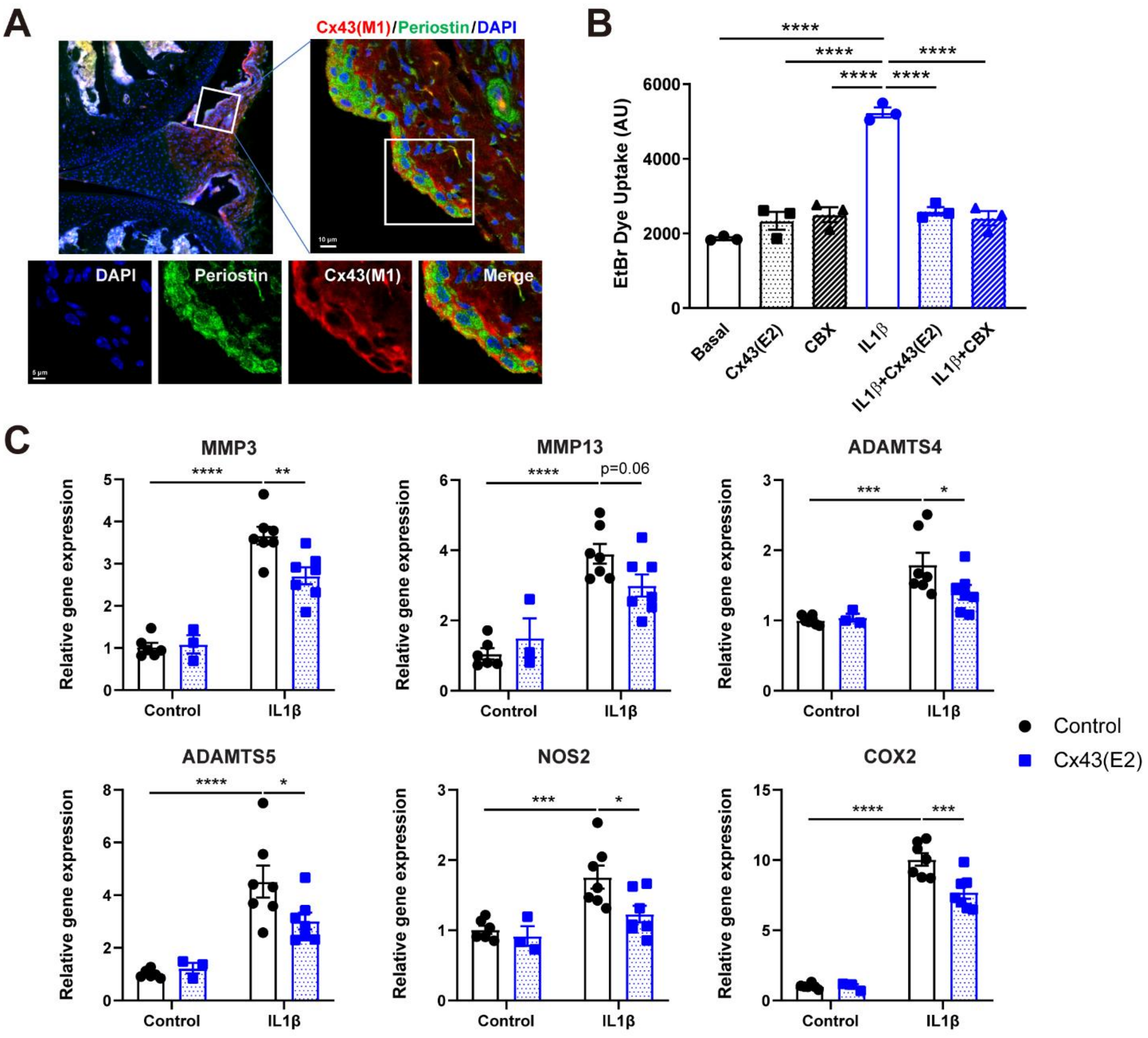


**Figure S6. Cx43 hemichannel-blocking antibody targeted synovial fibroblasts and decreased inflammatory gene expression.**

**(A)** Tissue sections were immunolabeled for the fibroblast marker Periostin, with one synovial region shown at higher magnification in the lower panels. **(B)** Human synovial fibroblast cell line SW982 was treated with IL1β with or withoutCx43(E2) antibody or CBX. The level of EtBr dye uptake was quantified by fluorescence microcopy and NIH ImageJ. **(C)** RNA was extracted from SW982 cells treated with IL1β and Cx43(E2) antibody and subjected to qRT-PCR analysis for MMP3, MMP13, ADAMTS4, ADAMTS5, NOS2, and COX2 mRNA expression. n =3-7/group. Data are presented as mean ± SEM. *, $p < 0.05$; **, $p < 0.01$; ***, $p < 0.001$; ****, $p < 0.0001$.

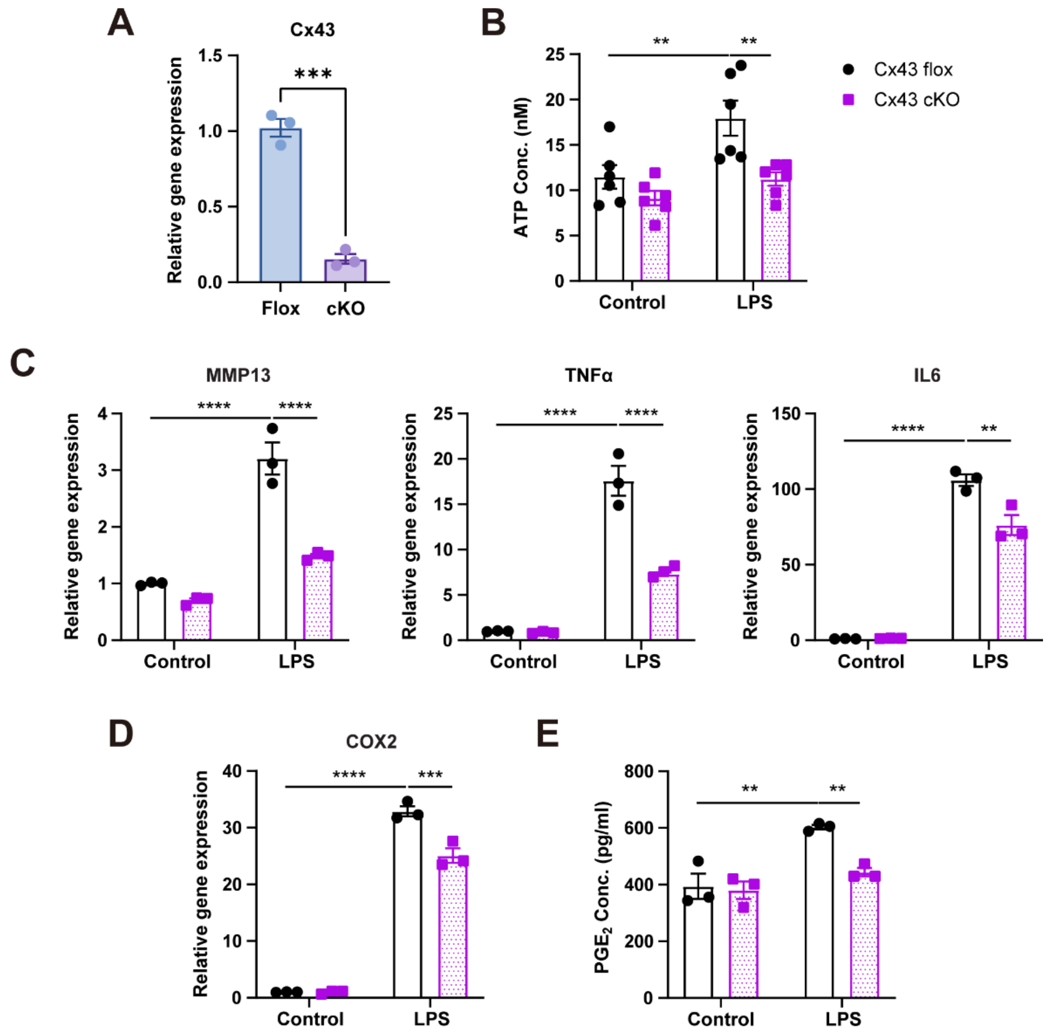


**Figure S7. Cx43 deletion in peritoneal macrophages reduced LPS-induced inflammatory gene expression, accompanied by decreased ATP and $PGE_2$ release.**

**(A)** Peritoneal macrophages were isolated from Cx43 flox/flox or Cx43 cKO mice and analyzed for Cx43 expression. n = 3 mice/group. **(B)** Extracellular ATP release was determined in peritoneal macrophages treated with LPS and Cx43(E2) antibody. **(C-D)** RNA was extracted and subjected to qRT-PCR analysis for MMP13, TNFα, IL6, and COX2 mRNA expression. **(E)** Extracellular $PGE_2$ release was determined in peritoneal macrophages treated with LPS and Cx43(E2) antibody. n =3-6/group. Data are presented as mean ± SEM. **, $p < 0.01$; ***, $p < 0.001$; ****, $p < 0.0001$.

**Table S1 Primers sequences for RT-PCR**

| Gene | Forward primer (5'-3') | Reverse primer (5'-3') |
|---|---|---|
| Human MMP3 | GTGAGGACACCAGCATGAA | GACCACTGTCCTTTCTCCTAAC |
| Human MMP13 | TCCTGATGTGGGTGAATACAATG | GCCATCGTGAAGTCTGGTAAAAT |
| Human ADAMTS4 | CTATGGGCACTGTCTCTTAGACAAAC | CACTGGCGGTCAGCATCA |
| Human COX2 | CTCAGCCATACAGCAAATCCT | CCGGGTACAATCGCACTTAT |
| Human IL6 | AAAGAGGCACTGGCAGAAA | CAGGCAAGTCTCCTCATTGAA |
| Mouse Cx43 | GCCGGCTTCACTTTCATTAAG | GAGTAGGCTTGGACCTTGTC |
| Mouse MMP3 | GGACCAGGGATTAATGGAGATG | TGAGCAGCAACCAGGAATAG |
| Mouse MMP13 | CCCTGATGTTTCCCATCTATACC | TTCATCGCCTGGACCATAAAG |
| Mouse Adamts4 | AGACGAAGCACTCACCTTGG | CTCCAGCCTGAGGAACATTGA |
| Mouse Adamts5 | AAATGGCAGCACCAACATAAC | CCAGGGTGTCACATGAATGA |
| Mouse TNFα | CATCTTCTCAAAATTCGAGTGACAA | TGGGAGTAGACAAGGTACAACCC |
| Mouse IL6 | CCAGAGTCCTTCAGAGAGATACA | AATTGGATGGTCTTGGTCCTTAG |
| Mouse COX2 | CGGACTGGATTCTATGGTGAAA | CTTGAAGTGGGTCAGGATGTAG |
| Mouse NOS2 | GGAATCTTGGAGCGAGTTGT | CCTCTTGTCTTTGACCCAGTAG |
